# Transcriptional signatures in the rat medial prefrontal cortex associated with vulnerability and resilience across distinct phases of opioid use disorder

**DOI:** 10.1101/2024.02.29.582799

**Authors:** Shirelle X. Liu, Peter Muelken, Zia L. Maxim, Aarthi Ramakrishnan, Molly S. Estill, Mark G. LeSage, John R. Smethells, Li Shen, Phu V. Tran, Andrew C. Harris, Jonathan C. Gewirtz

## Abstract

We characterized gene transcriptional activity in the medial prefrontal cortex of rats associated with individual differences in vulnerability to three distinct phases of opioid use disorder (OUD). Resilient rats showed many more changes in canonical pathway activity than Vulnerable rats in models of both early and advanced OUD, involving passive opioid exposure and opioid self-administration (SA), respectively. The Resilient/Vulnerable phenotype was also associated across phases with functionally specific gene networks, including those mediating epigenetic, neuroimmune, and neuroplasticity function. In contrast, we identified two phase-specific effects. First, differential activity of a myelination-related gene network was associated with Resilience/Vulnerability measured after passive morphine exposure. Second, expression of the calmodulin-inhibitor *Pcp4,* a gene recently implicated in a rat opioid SA GWAS analysis, was associated with Resilience/Vulnerability measured after SA but not after passive morphine exposure. Thus, we have identified both general and phase-specific transcriptional signatures involved in OUD vulnerability across its trajectory.

**Teaser:** Adaptations in the brain transcriptome are associated with resilience and vulnerability to opioid use disorder.

## Introduction

With a dramatic rise in the misuse of opioids, opioid use disorder (OUD) imposes a substantial and growing health burden in the US and globally (*1*, *2*). While this highlights the addictive potency of opioids (*3*), individuals still show substantial variability in their susceptibility to OUD due to genetic and environmental factors (*4*). Developing novel strategies for prevention and treatment of OUD requires a better understanding of the biomolecular mechanisms underlying the disease’s trajectory from initial opioid exposure to compulsive and relapsing opioid use.

Like substance use disorders in general, the chronic and relapsing nature of OUD suggests the influence of stable, epigenetic changes in gene regulation (*5*, *6*). While studies investigating effects of opioids on gene expression in rodent behavioral models have traditionally focused on differences between groups (i.e., opioid-versus saline-exposed animals), recent research has begun to elucidate patterns of gene transcription underlying individual differences in the severity of OUD-like behavior in rodents after prolonged opioid self-administration (SA) (*7*, *8*). However, epigenetic markers of vulnerability may arise earlier in the trajectory of OUD, especially considering the rapid transition from initial to compulsive opioid use (*3*). This possibility is also suggested by evidence that the early, acute behavioral responses to opioids and other addictive drugs predict the severity of later vulnerability to addiction (*9–17*), and by the overlap between transcriptional signatures of acute heroin exposure and chronic heroin SA (*7*). Thus, the goal of the current study was to characterize transcriptional mechanisms of vulnerability to OUD, from early opioid exposure to advanced OUD.

To characterize vulnerability to the development of OUD induced by initial opioid exposure (i.e., “early” OUD), we categorized rats based on their levels of withdrawal-induced anhedonia (WIA), as indexed by the degree of elevation in intracranial self-stimulation (ICSS) thresholds during withdrawal from acute, experimenter-administered morphine (*18–20*). This measure is unique in that low WIA predicts high levels of multiple indices of subsequent morphine SA and therefore provides a prospective behavioral model of vulnerability to OUD (*20*). The use of this passive-exposure paradigm also avoids several factors intrinsic to SA models (e.g., differences in drug intake and days of exposure between animals) that can confound interpretation of molecular data.

To characterize transcriptional mechanisms associated with individual differences in the advanced stages of OUD, we utilized two measures of drug SA that mirror different diagnostic features of the disorder. First, we used the behavioral economic measure of Demand to assess the persistence of drug-taking in the face of increased behavioral “cost”. This measure has been validated across species, including in human studies of several substance use disorders including OUD (*21*, *22*). Second, we measured transcriptional patterns associated with the propensity to relapse to opioid seeking using a cue/drug-induced reinstatement paradigm, the neurobiological substrates of which have been extensively investigated (*23*).

We focused our molecular analysis on gene transcription in the medial prefrontal cortex (mPFC), which is critically involved in all phases of addiction in both humans and rodents (*24–28*). Our findings revealed a variety of transcriptional adaptations associated with one or more behavioral paradigms, and with either drug treatment (i.e., morphine versus saline) or explicitly with the trait of OUD vulnerability (i.e., vulnerability versus resilience). In both our pre-emergence (i.e., WIA) and relapse (i.e., Reinstatement) behavioral phenotypes of Resilience/Vulnerability to OUD, most of the latter adaptations were specific to resilient rats and thus may serve a protective function. As such, our findings provide guideposts for the discovery of new therapeutic approaches to reduce the development of OUD and to tailor treatments to vulnerable individuals.

## Results

### Individual differences in variability in three behavioral measures that model the trajectory of OUD

Fig. 1 shows an overview of our experimental design. We first tested rats in one of three behavioral paradigms – WIA, Demand, or Reinstatement – to capture OUD’s stages of initial acute opioid exposure, persistent opioid use, and relapse, respectively.

**Fig. 1.**
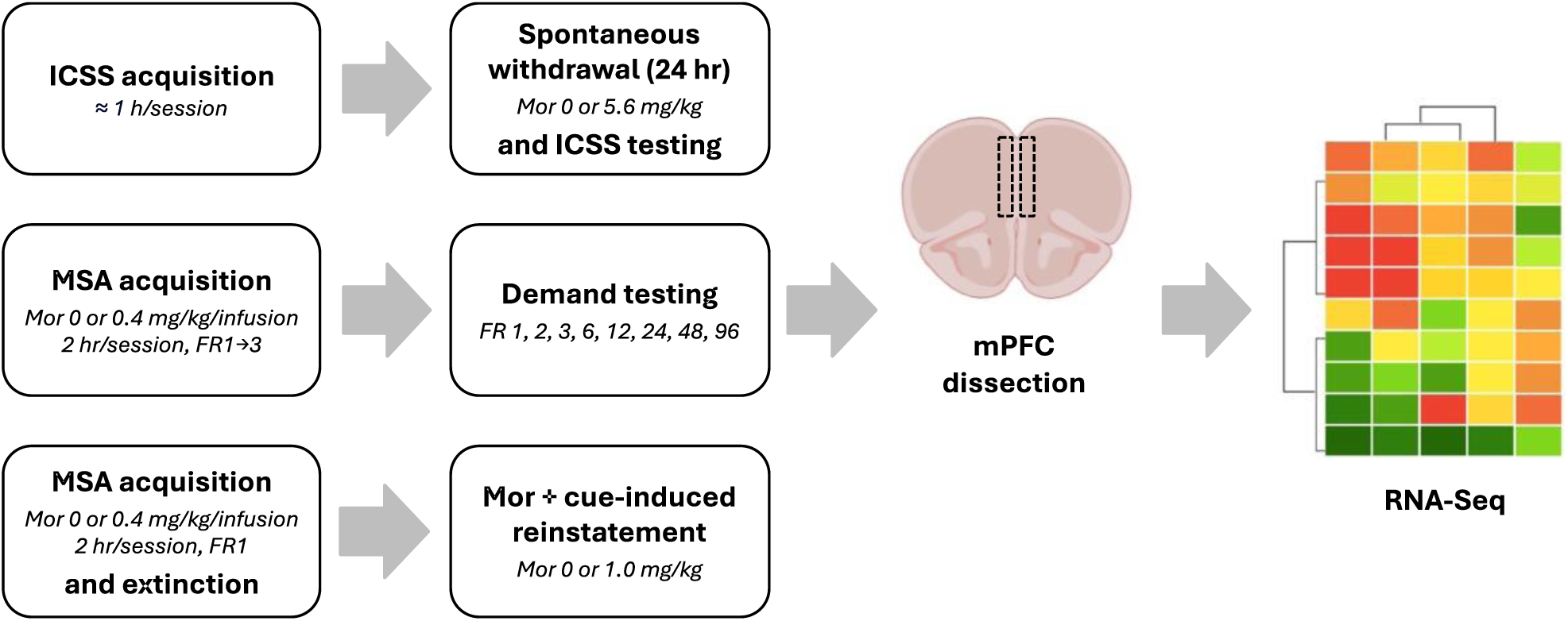
Experimental design. Timeline of the three behavioral paradigms, mPFC collection, and RNA-seq analysis is shown.

To measure WIA, rats received repeated daily injections of either morphine sulfate (5.6 mg/kg, s.c. n = 26, 10-16/sex) or saline (n = 13, 5-8/sex) and ICSS thresholds were tested 23 h after each injection (i.e., during spontaneous withdrawal) (*20*). Increases in ICSS thresholds represent a sensitive and reliable measure of anhedonia and have been observed during withdrawal from a range of addictive drugs (*20*, *29*). Groups did not differ in their baseline ICSS thresholds or response latencies (Table S1). A three-way ANOVA indicated a significant effect of group (*F* (1, 35) = 20.3, *p* < 0.0001), with ICSS thresholds significantly elevated in morphine-compared to saline-treated rats (Fig. 2A). There were no significant effects of sex or session or interactions between these variables. Groups did not differ in their ICSS response latencies (Fig. S1A), indicating that elevations in ICSS thresholds were not attributable to non-specific (e.g., motoric) effects (*30*).

**Fig. 2.**
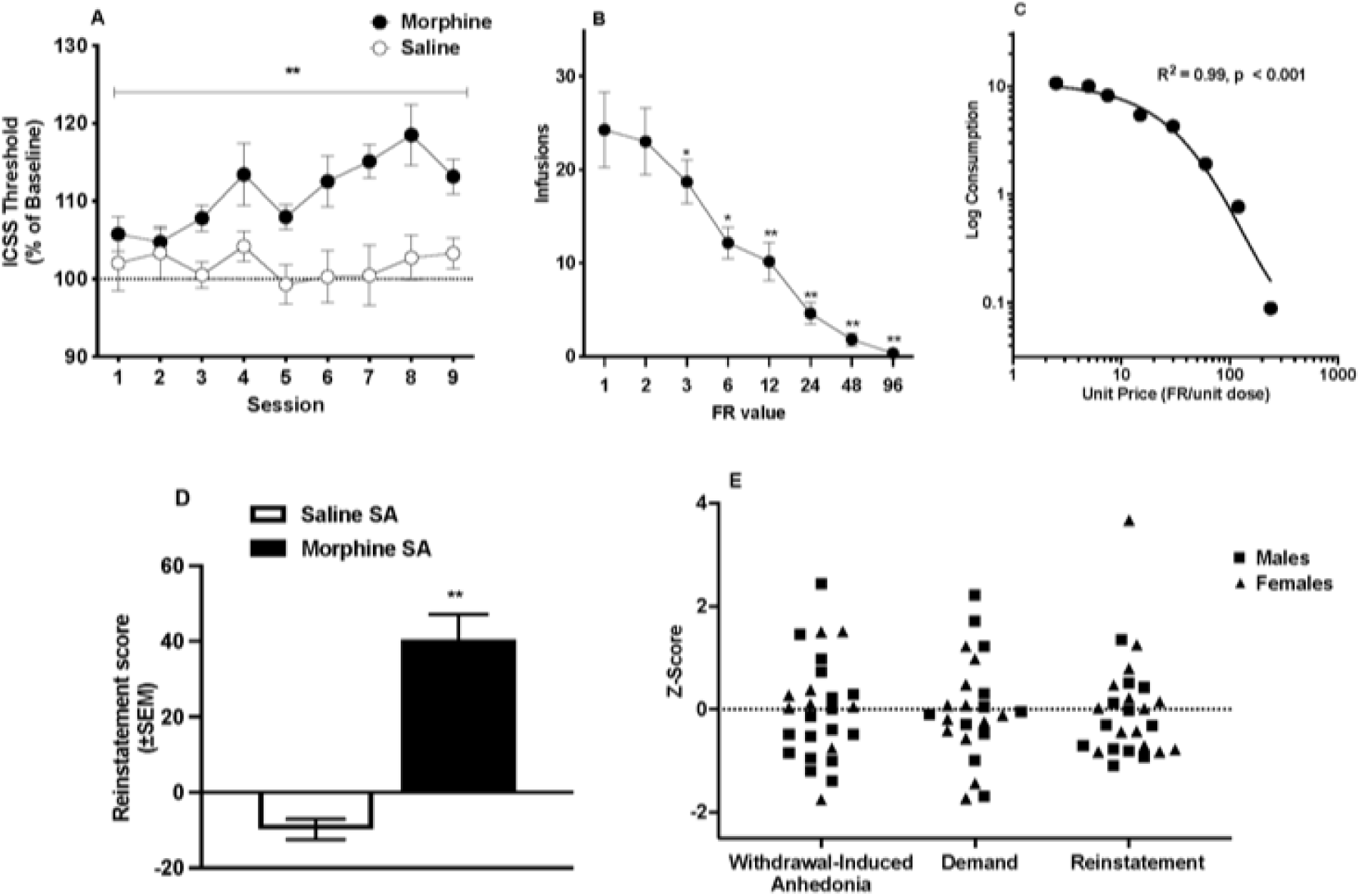
Male and female rats exhibit individual difference in vulnerability in three behavioral paradigms. **(A)** ICSS thresholds (% Baseline, Mean ± SEM) during spontaneous morphine withdrawal. ** Different from Saline (main effect), p < 0.0001. **(B)** Infusions at each FR (Mean ± SEM) during demand testing. *,** Different from FR 1, p < 0.05 or 0.01. **(C)** Exponential demand function for morphine during demand testing. **(D)** Reinstatement scores (Mean ± SEM) during the reinstatement test following morphine or saline SA. ** Different from saline SA, p < 0.01. Data are collapsed across sex in Figs 1A – 1D because there were no significant effects of sex in any behavioral model. **(E)** Z-score distribution for individual morphine rats of each sex is shown in panels (A)-(D). Data in (B) are a subset of animals from a larger cohort previously reported in Harris et al. (2024) (*93*).

To measure Demand, a separate group of rats trained to self-administer morphine (0.4 mg/kg/infusion) under a fixed ratio (FR) 3 schedule of reinforcement underwent a FR escalation protocol in which the response requirement was progressively increased each day (FR 1, 2, 3, 6, 12, 24, 48, and 96). Rats reliably acquired morphine SA, with the morphine group (n = 23, 11-12/sex) earning significantly more infusions than saline controls (n = 11, 5-6/sex) by the end of the acquisition period (Fig. S1B). Analysis of infusions in the morphine group during demand testing indicated a significant effect of FR (*F* (1.6, 32.3) = 29.4, *p* < 0.0001), reflecting a progressive reduction in infusions as the FR was increased, but no significant effects of sex or FR x sex interaction. When data were collapsed across sex, infusions decreased significantly at FR ≥ 3 compared to FR 1 (Dunnett *q* = 2.9 - 6.3, *p* < 0.05 or 0.01; Fig. 2B). Morphine consumption was well-described by an exponential demand function, with *R^2^* values typically ≥ 0.85 for individual animals (Table S2) and *R^2^* = 0.99 for rats as a group (Fig. 2C).

To measure reinstatement of morphine-seeking behavior, a separate group of rats trained to self-administer morphine (0.4 mg/kg/infusion) under an FR 1 schedule (n = 26, 12-14/sex) underwent saline extinction in the absence of the drug-associated cue light. Extinction of morphine SA resulted in a significant reduction in active lever presses (Fig. S2). To test reinstatement, rats were injected with morphine (1.0 mg/kg, s.c.) combined with presentation of the drug-associated light cue (i.e., drug + cue-induced reinstatement). A control group (n = 13, 6-7/sex) underwent Saline SA followed by “extinction” of SA and “reinstatement” induced by combined exposure to s.c. saline + the cue light. A two-way ANOVA on Reinstatement scores indicated a main effect of training drug (Morphine vs. Saline; *F* (1, 36) = 26.86, *p* < 0.0001; Fig. 2D), reflecting greater responding on the active lever in the morphine group on the test day (Fig. S2), but no main effect of sex or interaction with sex.

Given the substantial individual variability in all three behavioral measures (Fig. 2E),we were able to classify morphine-exposed rats as “resilient” and “vulnerable” in each paradigm using a median split of their behavioral scores. Since WIA is negatively correlated with subsequent morphine SA (*20*), resilient and vulnerable subgroups for WIA were those with high and low WIA, respectively. The behavioral economic measure α is inversely related to reinforcement efficacy (*31*). Resilient and vulnerable rats for Demand were therefore those associated with high and low α values, respectively. Resilient and vulnerable rats for Reinstatement were those associated with low and high reinstatement scores, respectively.

### Transcriptional signatures of Resilience and Vulnerability across the three paradigms

Twenty-four hours after the final drug exposure, we collected tissue from the mPFC of Morphine or Saline rats for RNA-seq (Fig. 1). This timepoint is commonly used in RNA-seq studies (*7*, *32*, *33*) and one at which we find substantial anhedonia during morphine withdrawal (*20*). We first conducted a Principal Component Analysis (PCA) of the 500 genes showing the highest expression levels. Drug treatment accounted for 15% –23% of the variation in male and female transcriptomes across the three paradigms, while sex accounted for 17% –25% of the variation (Fig. S3). Given the pronounced effects of sex, sequencing data were analyzed separately for each sex where appropriate.

We identified differentially expressed genes (DEGs) by comparing each of the resilient and vulnerable groups with their respective Saline controls within each paradigm (Tables S3 and S4). Contrary to our expectations, Ingenuity Pathway Analysis (IPA) in male rats revealed a far greater number of significantly altered biological pathways in resilient rats than in vulnerable rats in the WIA (*p* < 0.0001) and Reinstatement (*p*<0.0001) paradigms (Fig. 3A). The numbers of enriched canonical pathways in resilient and vulnerable male rats did not differ significantly in the Demand paradigm (*p* = 0.41). The top altered canonical pathways in males were associated with signaling pathways and plasticity (e.g., CREB, G-protein, cAMP signaling) and neuroimmune pathways (e.g., S100, phagosome, macrophage) (Fig. 3B). Consistent with the male data, we also observed a greater number of significantly altered biological pathways in resilient than in vulnerable female rats in the WIA (*p* = 0.017) and Reinstatement (*p* < 0.0001) paradigms (Fig. 3C). However, the opposite pattern was observed in females for Demand, with a greater number of canonical pathways altered in female vulnerable rats compared to female resilient rats (*p* < 0.0001). As observed in males, several of the top altered canonical pathways in females were associated with cell signaling and plasticity (e.g., G-protein) and neuroimmune function (e.g., phagosome, cytokine storm) (Fig. 3D). Taken together, these findings indicate that changes in canonical activity in resilient rats were greater than in vulnerable rats in the WIA and Reinstatement paradigms although not in Demand, with neuroimmune and plasticity pathways most consistently implicated across all three paradigms as a function of the Resilience/Vulnerability trait.

**Fig. 3.**
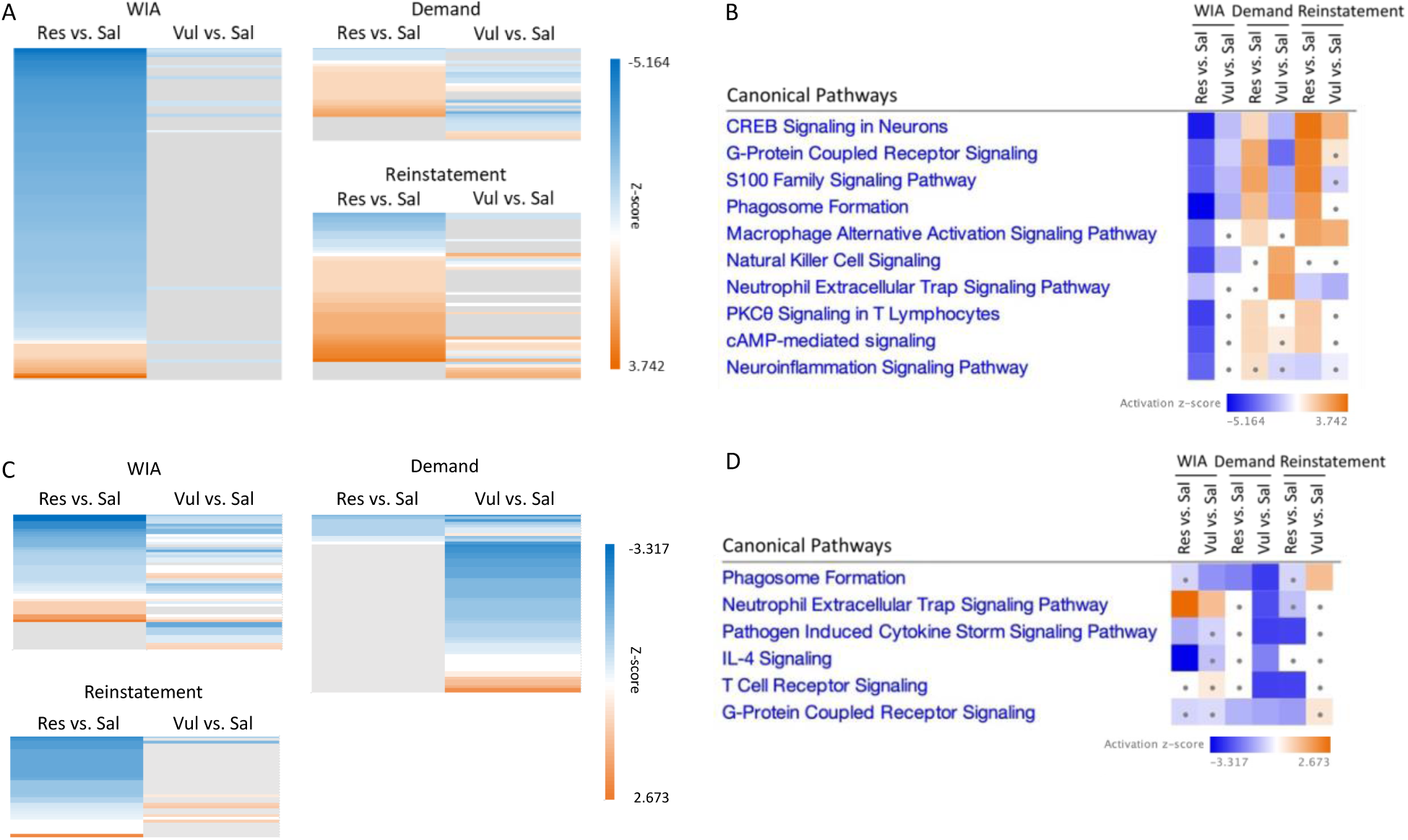
Resilient rats showed greater changes in IPA canonical pathways than vulnerable rats in WIA and Reinstatement. Heatmaps of all canonical pathways with detectable alteration in resilient versus saline and/or vulnerable versus saline groups in each behavioral paradigm in male (A) and female (C) rats. Each line represents one pathway. Canonical pathways serving neuroimmune- or plasticity-related functions within the top 15 canonical pathways ranked by z-score in male (B) and female (D) rats. Grey, z-score = N/A; Square with a dot, absolute z-score < 1.

Next, we conducted a variance partitioning analysis (*34*) to identify drivers of variation in gene expression attributable to Resilience/Vulnerability common to all three behavioral paradigms. Despite substantial procedural differences between the paradigms (e.g., patterns, dose, and contingency of morphine exposure, etc.), the Resilience/Vulnerability trait accounted for up to 32% and 36% of the variation in expression of individual genes in male and female morphine animals, respectively (Figs. 4A-D). Gene ontology (GO) analysis of the 754 genes in males where the Resilience/Vulnerability trait accounted for at least 10% of variation in expression values showed enrichment of functions associated with epigenetic regulation, including export of proteins from the nucleus, chromatin remodeling, and nucleic acid metabolic processes (Fig. 4E). GO analysis of the 372 genes meeting the same criterion in females showed enrichment of genes associated with cell signaling, including GTPase cycle and signal transduction functions (Fig. 4F).

**Fig. 4.**
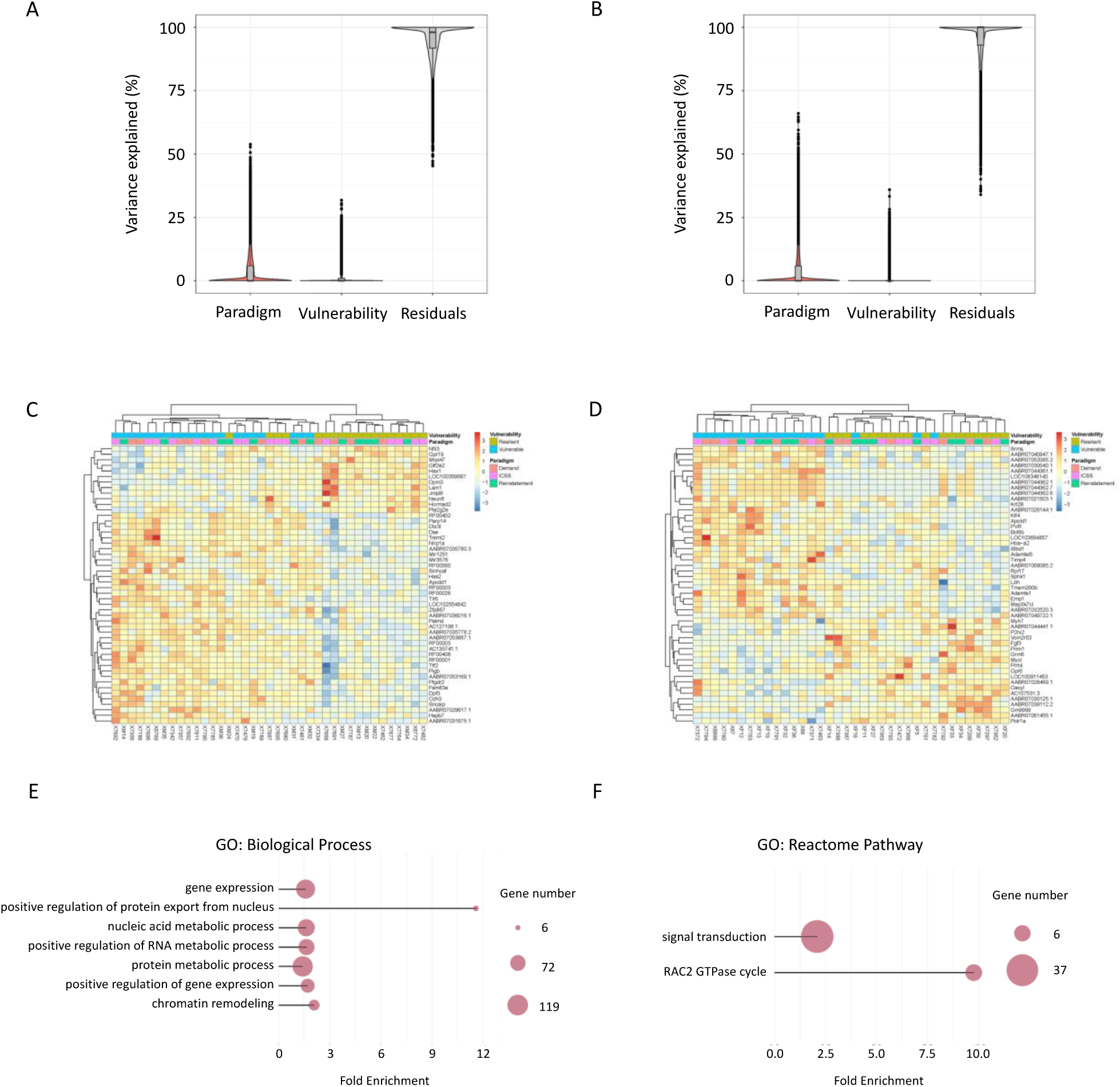
Variations in gene expression attributable to vulnerability across the 3 behavioral paradigms. (A, **B)** Variance partitioning analysis yielded the percentage of variance explained by paradigm, vulnerability, or residuals in male **(A)** and female **(B)** rats. **(C, D)** Heatmaps of the 50 genes in male **(C)** and female **(D)** rats where vulnerability accounted for the greatest proportion of variance in expression levels. **(E)** Genes where vulnerability accounted for >10% of variance were associated with epigenetic regulation in male rats. **(F)** Genes where vulnerability accounted for >10% of variance were associated with cell signaling and the RAC2 GTPase cycle in female rats.

### Transcriptional Signatures of Early or Late OUD

To identify functional gene networks and their putative regulators altered by morphine exposure, we conducted Weighted Gene Co-Expression Network Analysis (WGCNA) (*35*) to build co-expression modules based on gene activity within each of our three behavioral paradigms. This was conducted on rats collapsed across sexes and the Resilience/Vulnerability trait to maximize network reliability (*36*). In the WIA paradigm, differential gene co-expression between morphine- and saline-treated rats was detected in five WGCNA modules. Gene Ontology (GO) analysis identified functional specificity in only one module (“saddlebrown”), which showed reduced gene co-expression in morphine-treated rats (MeDC = -0.38, *p* = 0.02). Gene function in this module was strongly associated with oligodendrocyte maturation and myelination (Fig. 5A). Algorithm for the Reconstruction of Gene Regulatory Networks (ARACNe) (*37*) revealed the regulatory effects of two transcription factors (TFs), PRDM8 and HR within this module (Fig. 5B). Both TFs are epigenetic histone modifiers, with PRDM8 functioning as a histone methyltransferase and HR as a histone lysine demethylase. Notably, PRDM8 plays a role in regulating oligodendrocyte differentiation (*38*).

**Fig. 5.**
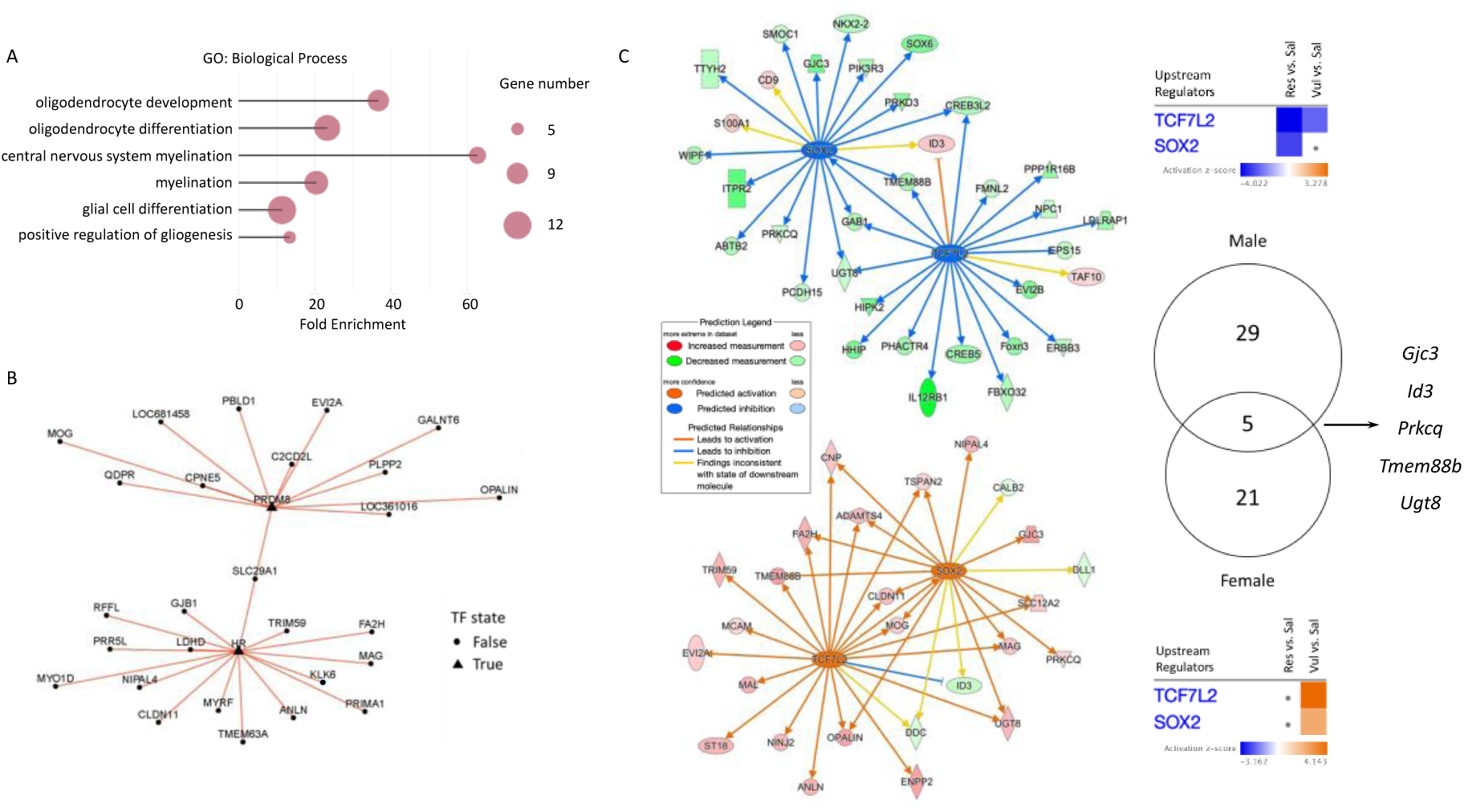
Myelination/oligodendrocyte processes exhibited weaker activities in resilient groups in WIA rats in both sexes with distinct molecular mechanisms. **(A)** A WGCNA “saddlebrown” module with decreased connectivity in Morphine rats showed enrichment of genes functioning in oligodendrocyte processes and myelination. **(B)** ARACNe analyses of genes in the saddlebrown module implicated the transcription factors (TFs) PRDM8 and HR. **(C)** IPA upstream regulator analysis predicted the activity of TCF7L2 and SOX2 with more inhibition in resilient WIA males, and with more activation in vulnerable WIA females. Square with a dot, absolute z-score < 1. Venn diagram showing 5 overlapping genes between WIA resilient male and vulnerable female TCF7L2 and SOX2 networks.

To further investigate whether gene network activity was altered in relation to the Resilience/Vulnerability trait for WIA, we conducted an IPA analysis to identify upstream regulators of genes that were differentially expressed in Vulnerable or Resilient rats compared to Saline controls. This analysis predicted altered regulation of two TFs, TCF7L2 and SOX2, in both sexes (Fig. 5C, top panel, male; bottom panel, female). Both TFs play critical roles in myelination (*39–41*). The effects in the two sexes were consistent in directionality, in that the two TFs were predicted to be more strongly inhibited in the resilient group in males and more strongly activated in the vulnerable group in females. Even though male and female networks, comprising 34 and 26 DEGs, respectively, shared only five common DEGs, it is notable that activation of these TFs was associated with greater OUD vulnerability in both sexes (Fig. 5C).

We next conducted WGCNA analyses on the advanced OUD paradigms of Demand and Reinstatement. Differences in gene co-expression between morphine- and saline-treated rats were detected in six WGCNA modules in the Demand paradigm and six modules in the Reinstatement paradigm. While none of the Reinstatement modules exhibited functional specificity, one of the Demand modules (“White”) showed significant enrichment of genes associated with various aspects of neuroimmune function (MeDC = - 3.81, *p* < 2.20E-16) (Fig. S4), consistent with the canonical pathway analysis described above (Fig. 3B and 3D).

### Coherence with other rat and with human datasets

A recent genome-wide association study (GWAS) on Heterogenous Stock (HS) rats revealed genomic loci associated with behavioral measures of heroin exposure and SA (*42*). The HS outbred strain is particularly well-suited for genomic analyses, with high genetic diversity and the availability of complementary datasets such as whole-genome sequencing and multi-tissue-type expression quantitative trait loci (eQTL) data. Accordingly, Kuhn et al. (2025) revealed significant causal coding variants in three genes that were in strong linkage disequilibrium (LD) with a region on chromosome 11 associated with heroin consumption. The same region was also in high LD with eQTLs for expression of two of these genes – SH3 domain binding glutamate-rich protein (*Sh3bgr*), and Purkinje cell protein 4 (*Pcp4)* – in the mPFC. To further investigate the potential functional relevance of these genes to individual differences in OUD, we asked whether their expression in the mPFC varied systematically as a function of our three behavioral indices of Resilience/Vulnerability. Consistent with the GWAS and eQTL data in HS rats, the differential slope of *Pcp4* expression varied significantly in males as a function of Demand (log2FC = 0.27, *p* < 0.01; *R^2^* = 0.23, *p* = 0.13) and Reinstatement (log2FC = - 0.25, p < 0.01; *R^2^* = 0.49, *p* = 0.01), although the direction of the relationship (positive versus negative) differed between the two behavioral measures (Fig 6B, C). The differential slope of *Pcp4* expression did not vary significantly as a function of in WIA in males (Fig. 6A). The slope functions for *Pcp4* in female rats, or for *Sh3bgr* in either sex, were not significant (Fig. S5).

**Fig. 6.**
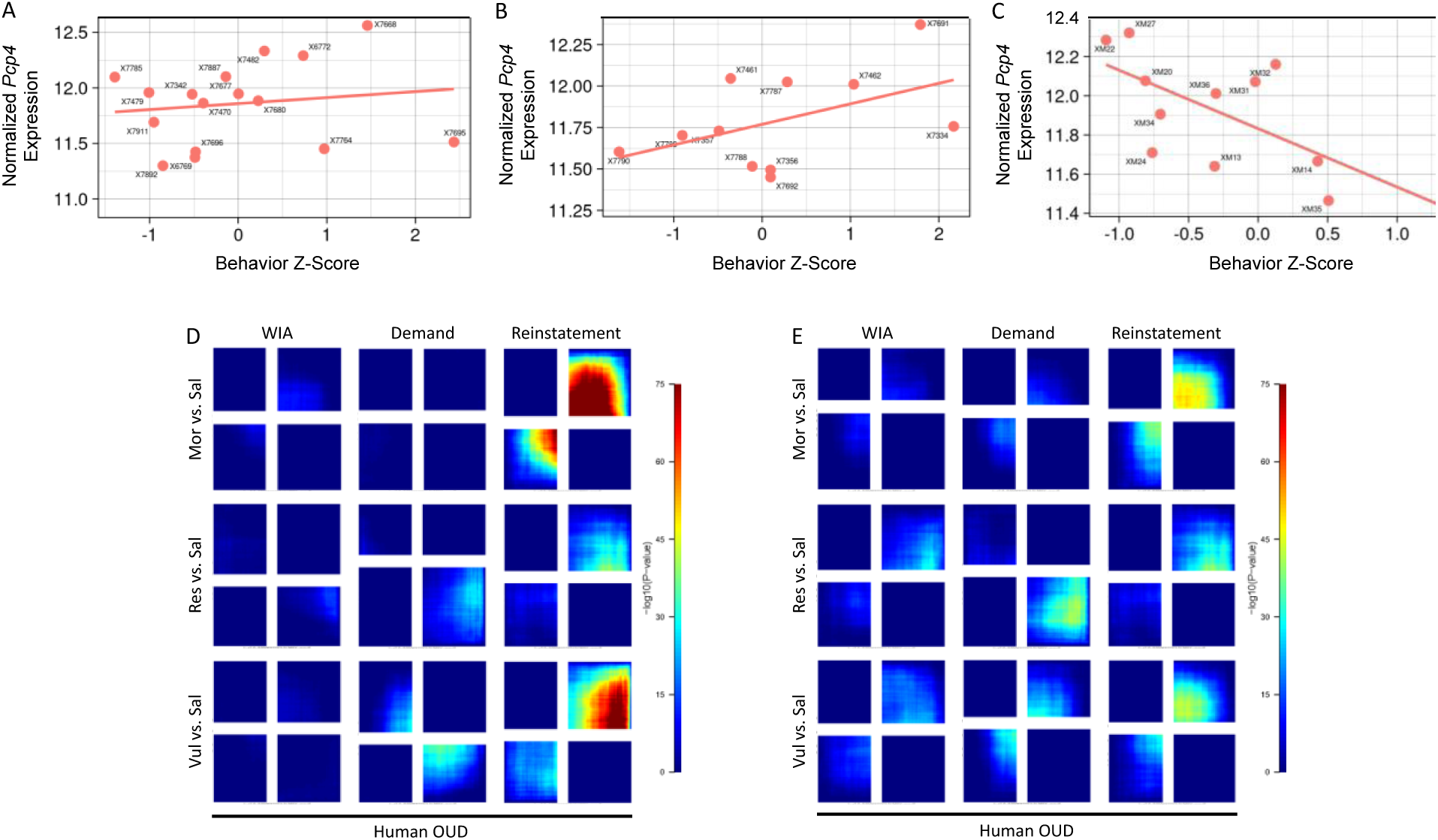
Coherence with other rat and with human datasets. (A-C) Correlation between normalized *Pcp4* expression and behavioral Z-Scores in male WIA **(A)**, Demand **(B)**, and Reinstatement **(C)** rats. **(D, E)** RRHO comparing gene expression patterns in the NAc of human OUD patients vs. controls and morphine-exposed, resilient, or vulnerable rats vs. control rats in males **(D)** and females **(E)** in the 3 paradigms.

To evaluate the translational relevance of our measures of vulnerability across the three paradigms, we next assessed shared patterns of gene expression in the mPFC in our rat behavioral models and in the dorsolateral prefrontal PFC (dlPFC) and nucleus accumbens (NAc) of human OUD patients postmortem (*43*). The threshold-free approach of Rank-Rank Hypergeometric Overlap (RRHO) revealed pronounced overlap between mPFC transcriptional expression in rats vulnerable to Reinstatement and in the dlPFC and NAc of both male and female OUD patients. The magnitude of this effect was stronger in the NAc (Figs. 6D, 6E, S6), consistent with its higher functional connectivity with the mPFC (*44*, *45*). The overlap effects in the NAc were primarily driven by vulnerable animals in both male (Fig. 6D) and female groups (Fig. 6E). In contrast, there was only modest overlap (Demand) or little or no overlap (WIA) between transcriptional expression in males and females in the other behavioral paradigms and OUD patients (Figs. 6D, E).

## Discussion

The current study identified patterns of gene network activity in the rat mPFC associated with vulnerability to OUD at different stages across the disorder’s trajectory. In the early (i.e., prospective) and in one of the advanced (i.e., relapse) behavioral models of OUD, our analyses revealed greater transcriptional reprogramming in rats that were resilient to OUD than in vulnerable rats. This core trait of Resilience/Vulnerability was associated with alterations in activity of gene networks involved in epigenetic and neuroimmune function and in signal transduction. In contrast to these multi-stage findings, we also identified a myelination-related gene network in which gene expression was associated specifically with Resilience/Vulnerability in our WIA paradigm involving experimenter-administered morphine. Conversely, expression of *Pcp4*, a gene recently implicated in a GWAS of heroin SA in rats, was linked to Resilience/Vulnerability in both of our SA paradigms (i.e., Demand and Reinstatement) but not in WIA. There was also marked overlap between the profile of gene expression specifically in Reinstatement and gene expression in human OUD overdose patients. Finally, while some of these effects were seen in both sexes, others were sex specific. By investigating transcriptional adaptions associated with the Resilience/Vulnerability phenotype both prospectively and in advanced phases of OUD, these findings identify distinct potential targets for novel therapeutics either to blunt the addictive potency of opioids in nonaddicted individuals (e.g., when administered opioids as anesthetics) or to treat those already diagnosed with OUD.

### The transcriptomics of Resilience/Vulnerability across the trajectory of OUD

A striking – and surprising – finding was the larger number of enriched canonical pathways in both male and female resilient rats compared to vulnerable rats in WIA and Reinstatement. Contrary to the common assumption that greater vulnerability to OUD largely reflects higher sensitivity to the effects of opioids (*46–49*), this suggests that vulnerability to OUD might also reflect a failure to mount appropriate adaptive responses to opioids (*50*). Furthermore, the fact that WIA and Reinstatement differ from each other in multiple ways (e.g., drug dose, experimental duration and contingency of opioid exposure) suggests that this difference in transcriptional responsiveness reflects a general feature of the Resilience/Vulnerability trait rather than a defining feature of a given paradigm. Further investigation and confirmation of the transcriptional pathways that are recruited in resilient individuals may thus yield novel approaches to mitigating susceptibility to OUD and/or promotion of recovery by increasing protective regulatory mechanisms.

The canonical pathways that were most affected by the Resilience/Vulnerability trait in all three paradigms were associated broadly with neuroimmune activity (e.g., phagosome formation) and activity-dependent neuroplasticity (e.g., CREB- and cAMP-dependent signaling). A neuroimmune-related functional module was also implicated in Demand in our WGCNA analysis. These findings complement a rapidly growing literature implicating neuroimmune activity in addiction to opioids and other drugs (*51–53*), as well as decades of research implicating cAMP/CREB-mediated neuroplasticity in addiction (*54*, *55*).

To further investigate the transcriptional machinery associated with the core trait of OUD Resilience/Vulnerability, we conducted a variance partitioning analysis (*34*) to parse out the variance in gene expression attributable to individual differences in Resilience/Vulnerability versus the variance attributable to our behavioral paradigms. This analysis revealed enrichment in males of gene pathways contributing to epigenetic regulation, such as chromatin remodeling and nuclear transport. Nuclear transport proteins have recently emerged as master regulators of genome organization through their shuttling of TFs between nucleus and soma (*56*, *57*). For example, two of the enriched genes in this category code for Exportin 4 (XPO4) and RanGTPase (RAN), proteins that form a complex to transport a number of transcription factors in and out of the nucleus through nuclear pores. These proteins also export circular RNAs, regulatory molecules that have been implicated in several studies in substance use disorders, including OUD (*56*, *58*).

While gross dysfunction of nuclear transport is linked to neurodevelopmental diseases and various forms of cancer (*59*, *60*), our findings suggest that more fine-tuned variations in their expression may play a role in vulnerability to OUD. In females, variance partitioning analysis showed enrichment of genes associated with associated with the RAC2 GTPase cycle, which have been implicated in addiction to opioids and other drugs (*61*, *62*), as well as with signal transduction. Indeed, one gene in this family that was enriched, *Rasal1*, is necessary for neuronal maturation and promotes NMDA receptor-mediated synaptic transmission (*63*). Overall, our findings support the idea that variability in activity of gene networks associated with epigenetic regulation, neuroimmune function and neuroplasticity contribute to individual differences in vulnerability to OUD.

### Functional gene networks associated with individual differences in vulnerability in different phases of OUD

While the above findings indicate recruitment of common canonical pathways across the trajectory of OUD, other gene networks were implicated in a stage-specific manner. In the WIA paradigm, noncontingent opioid injections in previously naive rats altered connectivity in a network of genes associated with maturation of oligodendrocytes and consequent myelination. The TF-coding *Prdm8*, which regulates several other genes in the network, is a member of the PRDM family of TFs that broadly affect gene activity through chromatin regulation and are themselves regulated through DNA methylation (*64*). PRDM8 itself is specifically involved in repressing the production of oligodendrocyte precursor cells (*38*). Additionally, several of the functionally relevant genes in this network (*Mal*, *Mog*, *Plp*, and *Opalin*) are involved in the later stages of oligodendrocyte differentiation and myelination (*65*).

In order to maximize statistical power (*36*), this analysis did not differentiate between male and female or between vulnerable and resilient animals. To evaluate these factors, we compared sets of DEGs in each of the male and female vulnerable and resilient groups with DEGs in their respective saline control groups. Consistent with our network analysis findings, these analyses implicated two TFs, TCF7L2 and SOX2, in differential gene activity in vulnerable and resilient rats. These TFs positively regulate oligodendrocyte proliferation and differentiation, as well as myelination and remyelination post injury (*39*– *41*, *66*). Interestingly, the same two TFs were implicated as upstream regulators of gene networks in both males and females, even though the networks in each sex showed little overlap of downstream DEGs. Furthermore, whereas TCF7L2 and SOX2 were downregulated in resilient male rats, they were upregulated in vulnerable female rats. Thus, the net effect of both genes was the same in the two sexes.

While the participation of astrocytes in OUD in preclinical models is now well established (*67–69*), there has been less focus in the past on the role of oligodendrocyte proliferation. This has begun to change with a recent report that morphine promotes oligodendrogenesis in the mouse ventral tegmental area (VTA) and, conversely, that blockade of oligodendrogenesis in the VTA reduces dopamine release in the nucleus accumbens and induction of a morphine conditioned place preference (*70*). Oligodendrocyte precursor cells constitute the primary proliferative cell type in the adult CNS, receive neuronal synaptic inputs, show environmentally induced expression of glutamate receptors, and stabilize neuroplasticity (*71–73*). In this context, the current results suggest that stimulation of oligodendrocyte differentiation and myelination during withdrawal from morphine exposure may be a key player in the neuroplastic adaptations underlying the development of OUD.

### Putative role of PCP4 peptide in OUD vulnerability

The current study addressed epigenetic, but not genetic, mechanisms underlying vulnerability and resilience to OUD. However, our findings implicating *Pcp4* complement a recent GWAS study on heroin self-administration in HS rats (*42*), as well as evidence implicating *Pcp4* in nicotine and alcohol dependence in humans (*74*, *75*). Specifically, we found that levels of *Pcp4* expression in the mPFC varied significantly as a function of the Resilience/Vulnerability indices in both advanced OUD paradigms (Demand and Reinstatement). Although the mechanism by which this calmodulin-inhibiting peptide might affect the etiology of OUD is currently unknown, an analog of PCP4 in zebrafish was recently found to interact with dopamine D2 receptor activity to modulate levels of motivated behavior (e.g., feeding) (*76*). Together, these findings offer a compelling rationale for further investigating the role of PCP4 in mediating vulnerability to OUD.

### Directionality of differential gene expression

We observed a common feature in two sets of analyses of Resilience/Vulnerability conducted across the three paradigms, whereby similar pathways or genes were differentially enriched but with reversed directionality. Thus, the same sets of canonical pathways that were upregulated in rats characterized as resilient in the Reinstatement paradigm were downregulated in rats characterized as resilient in the WIA paradigm. Similarly, *Pcp4* gene expression was a positive function of vulnerability as indexed in the Demand paradigm but a negative function in the Reinstatement paradigm. These findings reflect the likelihood that our gene transcription assays captured snapshots in time of bidirectional processes. There are multiple phenomena where the same molecular processes are inhibited or excited by opioids or other addictive drugs as a function of the time that has elapsed since the most recent exposure to the drug. For example, glutamatergic AMPA-mediated synaptic transmission is potentiated during cocaine abstinence and then depressed following drug re-exposure (i.e., relapse) (*77*). Furthermore, echoing the current findings, many of the same genes that were upregulated in the orbitofrontal cortex 2 days after methamphetamine exposure were downregulated 35 days after methamphetamine exposure (*78*). To the extent that such bidirectional effects are a common motif of addiction-related molecular processes, future treatments may be more effectively directed towards stabilizing the activity of gene networks and molecular pathways rather than towards their simple excitation or inhibition.

### Translational significance

Rodent studies, such as the current one, are valuable in mapping the tissue-specific epigenetic landscape of disease pathology in parallel with human genomic studies. That said, inferences to the human condition are necessarily tentative, given substantial interspecies differences in the structure and function of the human and rodent nervous systems. One way to bridge this translational gap is to systematically compare transcriptional expression across the genome in humans with OUD and in animals that model the disorder. Along these lines, we found substantial overlap between gene expression patterns in the mPFC of rats vulnerable in our late-stage OUD paradigms (particularly Reinstatement) and gene expression patterns in the dlPFC and NAc of OUD patients (*43*). The convergence of transcriptional expression patterns in animal models of OUD and human patients seen here and elsewhere (*7*) supports the applicability of preclinical models for identifying molecular mechanisms underlying the compulsive and persistent use of opioids in human populations.

Although the magnitude of behavioral effects did not differ between males and females in any model of OUD, sex accounted for significant variation in the mPFC transcriptome and in multiple gene network analyses. This complements a report that male and female rodents exhibited similar incubation of morphine craving despite marked sex differences in the NAc transcriptome (*79*). Further characterization of the mechanism(s) accounting for sex differences in transcriptomic effects in models of OUD (e.g., role of sex hormones) could provide insights into sex-specific prevention and treatment strategies for OUD (*80*),.

In summary, this study reveals a number of trait-, sex, and stage-specific genes and gene networks that are associated with OUD as assessed through three rat behavioral models. As such, our findings highlight the importance of designing studies to explicitly parse the divergent effects of these three different dimensions on OUD vulnerability. Perhaps the most notable feature of our findings, however, is that several transcriptional mechanisms associated with the Resilience/Vulnerability phenotype were recapitulated in both sexes or in more than one behavioral paradigm (e.g., the same TFs were implicated in regulating gene expression related to this trait in both male and female rats, and Resilience was associated with a greater number of significantly altered canonical pathways in both WIA and Reinstatement paradigms). In this manner, the current findings suggest both individualized and general strategies and targets for reducing vulnerability to OUD across its trajectory.

## Materials and Methods

### Animals

Experimentally naïve male and female adult Sprague Dawley rats (Inotiv, Lafayette, IN) weighing 275-300 g (male) or 225-250 g (female) at arrival were used. All rats were individually housed in a temperature- and humidity-controlled colony room with unlimited access to water under a reversed 12-h light/dark cycle. Beginning one week following arrival, food was restricted to 16-18 g/day to facilitate operant performance, avoid detrimental health effects of long-term *ad libitum* feeding (e.g., obesity), and limit catheter migration in rats tested for morphine SA. All procedures were approved by the Institutional Animal Care and Use Committee (IACUC) of the Hennepin Health Research Institute and the University of Minnesota in accordance with the NIH Guide for the Care and Use of Laboratory Animals and the Guidelines for the Care and Use of Mammals in Neuroscience and Behavioral Research.

### Behavioral experiments

#### Withdrawal-induced anhedonia (WIA)

Rats were prepared and trained on a discrete-trial ICSS procedure (see Supplemental Material) in daily ≈ 1-hour sessions conducted Mon-Fri until ICSS thresholds were stable (<10% variability over 5 days). On the first test day, rats were tested for ICSS and then immediately injected with either 5.6 mg/kg morphine sulfate (n = 26, 16 males and 10 females; s.c.) or saline (n = 13, 8 males, 5 females). The morphine group was larger than the saline group in this and the other behavioral studies to allow stratification into high and low vulnerability groups (see below). ICSS was tested 23 hours later to measure spontaneous WIA, which was expected at this timepoint (*81–83*). All rats were then injected as on the preceding day. This injection regimen was repeated for 11 days, with ICSS tested 23 hours after each injection except for a 2-day weekend break in behavioral testing following the 5^th^ and 6^th^ injections. On these days, injections were administered at their usual time. Immediately after the final ICSS test (24 hours after the final morphine or saline injection), rats were sacrificed and tissue from multiple brain structures was harvested and preserved. This is a standard time point for evaluating effects of drugs or other manipulations on the transcriptome and epigenome (*84–87*).

#### Demand

A separate set of rats were implanted with i.v. catheters and allowed to respond for i.v. infusions of 0.4 mg/kg morphine (n = 23, 11 males, 12 females) or saline (n = 11, 5 males, 6 females) during daily 2-hr sessions conducted 7 days per week using our standard apparatus and procedures (see Supplementary Material). Rats were tested under a fixed ratio (FR) 1 schedule for at least 10 sessions and until robust responding was observed, at which point the FR response requirement was gradually increased to FR 3. After acquisition criteria were met (i.e., ≥ 10 sessions at FR 3 and ≥5 infusions per session, ≤20% coefficient of variation, and ≥2:1 response ratio on the active versus inactive lever across 3 sessions), demand was tested by increasing the FR requirement each day as follows: FR 1, 2, 3, 6, 12, 24, 48, 96. Morphine consumption under this protocol is well described by the current exponential demand function (*83*, *88*). Brains were harvested as described above, 24 hours after completion of Demand testing. Rats in the saline group did not acquire stable SA under the FR 3 schedule and therefore were not tested for demand. Brain collection in these animals occurred on the same day as that for a control-paired morphine SA rat that began the protocol at a similar time.

#### Reinstatement

A separate set of male and female rats (n = 26, 12 males, 14 females) were implanted with i.v. catheters and allowed to respond for morphine (0.4 mg/kg/infusion) under an FR 1 schedule as described above. Following at least 10 sessions and attainment of stable responding (same stability criteria as described for Demand), extinction conditions were introduced in which active lever responses resulted in saline infusions in the absence of the drug-associated cue light. Extinction was tested for a minimum of 21 sessions and until responding decreased by 75%, or 42 days of extinction, whichever occurred first. Reinstatement was then tested. During an initial “baseline” session, rats were given a saline injection (s.c.) 10 minutes prior to the session, and active lever responses continued to deliver saline infusions in the absence of the drug-associated cue light. To test morphine + cue-induced reinstatement, the following day rats were injected with morphine (1.0 mg/kg; s.c.) 10 min prior to the session. Active lever presses during that session resulted in saline infusions and the presentation of the drug-associated cue light. Brain samples were collected 24 hours later as described above. This morphine dose reliably elicits reinstatement (*83*). Reinstatement was induced by combined exposure to morphine + cue because this condition produces more robust reinstatement than either morphine or the cue alone (*83*). A negative control group (n = 13, 6 males, 7 females) was allowed to respond for i.v. infusions of saline throughout the protocol. To control for the visual stimulus conditions of the morphine group, saline infusions were delivered in the presence of the cue light for the same number of “acquisition” sessions as for a control-paired morphine SA rat. Likewise, the same number of “extinction” sessions were conducted as in the control-paired morphine SA rat and thus, no cue light was presented with saline deliveries. During “reinstatement”, rats were administered saline (s.c.) 10 min prior to both test sessions, wherein active lever presses had no consequences during the first “baseline” session but produced the saline-associated cue light during the second “reinstatement” test session. Brain samples in these rats were collected 24 hours later.

#### Statistical analysis

##### Withdrawal-induced anhedonia

ICSS thresholds and response latencies served as the measure of brain reinforcement function and general motoric function, respectively. Baseline ICSS thresholds (in µA) and response latencies (in sec) were defined as the mean during the last five sessions prior to withdrawal testing. ICSS thresholds and latencies during withdrawal testing were subsequently expressed as a percentage of baseline and analyzed using a 3-factor ANOVA with morphine dose and sex as between-subject factors and session as a within-subject factor. Rats were assigned to high and low WIA groups based on a median split of z-scores of average ICSS thresholds (% baseline) across all test sessions.

##### Demand

Infusions at each FR during demand testing in the morphine group were compared using a one-factor ANOVA with the FR requirement as a within-subject factor, followed by Dunnett’s post hoc tests comparing infusions at FR 1 to those at subsequent FRs. To measure demand, exponential demand-curve analyses were conducted as described in the Supplementary Material. The derived parameter α measures the degree to which a drug maintains SA following increases in unit price (FR / unit dose), and serves as an index of opioid reinforcement efficacy. Assignment to high and low demand groups was based on a median split of z-scores of α values.

##### Reinstatement

Assignment to high and low reinstatement groups was based on a median split of z-scores of reinstatement scores, defined using the formula: (active - inactive responses during reinstatement test) - (average of last 3 days of active - inactive responses during extinction). Differences in the reinstatement score was assessed using a 2-factor (sex x drug type) ANOVA, with a Šidák post-hoc test to determine within-factor differences.

### Brain tissue dissection

Rats were deeply anesthetized with isoflurane before decapitation. Brains were removed and dissected on an ice-cold metal block. Multiple brain regions from both hemispheres were isolated and aliquoted, flash-frozen in liquid nitrogen, and stored at -80°C. Only mPFC samples were further processed in the current study.

### RNA isolation, library preparation and sequencing

Total RNA was isolated using the RNAqueous™-Micro Kit (Invitrogen) according to the manufacturer’s protocol. Isolated RNA was quantified using the RiboGreen RNA Assay kit (Invitrogen) and assessed for quality using capillary electrophoresis (Agilent BioAnalyzer 2100; Agilent). Barcoded libraries were constructed for each sample using the TruSeq RNA v2 kit (Illumina). Libraries were assessed for quality using capillary electrophoresis (Agilent BioAnalyzer 2100; Agilent). RNA-seq libraries were size-selected for ∼200 bp fragments. Selected libraries were sequenced at a depth of ∼100 million reads per sample using NovaSeq 6000.

### RNA-seq analysis

Raw sequencing reads were aligned to the Ensembl v90 rat reference genome (rn6) using HISAT2 (v2.2.1) (*89*). Differential gene expression analysis was performed in R (v4.0.2) using the DESeq2 package (*90*). Significance was set at absolute log2(Fold Change) > 0.32 and *p* < 0.05 for RNA-seq (*32*).

### Weighted Gene Co-expression Network Analysis (WGCNA)

Sample expression values were normalized with variance stabilizing transformation (VST) using DESeq2. The biomaRt R package was used for extracting additional gene information, using the Ensembl Archive Release 104 (May 2021). The “removeBatchEffect” function from the limma package was used to remove the batch effect from the VST expression. WGCNA was used for module detection. In brief, the batch-adjusted VST expression was converted to an adjacency matrix using the “adjacency” function, with softPower=4. Topological overlap matrix similarity and dissimilarity was calculated using the “TOMsimilarity” function. Hierarchical clustering was performed on the adjacency matrix, using the “hclust” function (method=“average”). Requiring a minimum module size of 30 genes, the hierarchical clustering was cut into modules using the “cutreeDynamic” function. Numeric labels of modules were converted to colors using the “labels2colors” function. The eigengenes of the modules were calculated using the “moduleEigengenes” function, followed by a hierarchical clustering of the dissimilarity of the module eigengenes. Closely related modules were then merged using the “mergeCloseModules” function.

Once modules were generated, Differential Gene Connectivity Analysis (DGCA) was performed using the DGCA package in R, to determine the differential connectivity between morphine and saline conditions. Over-representation of differential gene sets with gene-associated modules was done using a hypergeometric test. Gene regulatory networks were calculated using ARACNE AP (version 2016-04-01) on batch-adjusted VST expression values, with 100 bootstrap iterations. Regulatory factors used in ARACNE were extracted from TRRUST (version2). Pearson correlation of module relationships to sample characteristics (i.e., “Behavior.Score”) was generated using the “cor” function. Network graphs were generated in R using the ggraph package.

### Ingenuity Pathway Analysis (IPA)

Differentially expressed genes (DEGs) were annotated by Ingenuity pathway Analysis (IPA; Qiagen) to identify relevant canonical pathways, molecular networks and cellular functions that showed significant alterations between comparison groups as previously described (*91*). Statistical significance (*p* < 0.05; -log(*p*) > 1.3) was determined by Fisher’s exact test. Fisher’s exact test was used to assess whether the number of significantly altered pathways (absolute Z-score ≥ 1) differed between the resilient versus saline and vulnerable versus saline comparisons.

### Variance Partitioning Analysis

“VariancePartition” package in R was used to compute the proportion of variance explained by paradigm and vulnerability in the data set (*34*). Counts for each sample were loaded into the R environment. For both male and female rats, we obtained the VST counts using DESeq2 for all genes. Batch effects were removed from the VST counts using the fitVarPartModel function. The formula for fitExtractVarPartModel was set to ∼ (1|Vulnerability) + (1|Paradigm) where Vulnerability takes on the values “Vulnerable” and “Resilient”, and Paradigm take on the values ICSS, Reinstatement and Demand. fitExtractVarPartModel function estimated the variation from Vulnerability differences for each gene across samples. Heatmaps sorted by the percent variance in genes explained by Vulnerability show the expression level of the top 50 most variable genes across the 2 vulnerability groups Resilient and Vulnerable as well as the 3 paradigms (Demand, WIA and Reinstatement). Heatmaps were generated using the R package “pheatmap”. Hierarchical clustering was applied for both rows and columns, and expression values were scaled for each row.

### Gene Ontology (GO) enrichment analysis

A list of genes from WGCNA or variance partitioning analyses served as the input for GO enrichment analysis (https://geneontology.org/). The reference list defines as all genes from the Rattus norvegicus database. The list of genes of interest was mapped to GO terms using the annotation functions “GO biological process complete” or “Reactome pathways”. The enrichment of each GO term was assessed using Fisher’s exact test. False discovery rate was used to adjust p-values for multiple comparisons.

### Gene expression - behavior score correlation analysis

For each paradigm - WIA, Reinstatement and Demand, morphine rats from 4 different batches were included in this analysis. Behavior scores indicating the vulnerability of rats to morphine addiction were scaled and centered for each paradigm. Differential analysis aimed to dissect the effect of behavior in male and female rats was performed after controlling for batch effects using a 2x2 factorial design with the following linear model, where the inclusion of the interaction term “Sex:Behavior.Score” facilitated in studying the Gender specific Behavior effect :

log(exp) ∼ Batch + Sex + Behavior.Score + Sex:Behavior.Score

This analysis facilitated in obtaining the genes associated with behavior scores in male and female rats, along with the slope (log2FoldChange) values indicating the strength and direction of association. The significance (padj) of the difference in the slope values were also obtained. The results are presented as a scatter plot between the behavior scores and the normalized expression obtained using DESeq2 for males and females separately using the R package ggplot2. Linear regression was used to calculate R² values for gene expression–behavior score correlations.

### Rank-rank hypergeometric overlap (RRHO)

The RRHO method was used to evaluate overlap in gene signatures between human OUD patients (assayed in dorsolateral PFC (dlPFC) and nucleus accumbens (NAc) postmortem by RNA-seq) and WIA, Demand, and Reinstatement data in rats in this study in a threshold-free manner (*92*). DEG files from human OUD samples were obtained from Seney et al. (2021) (*43*). Briefly, we ranked differential gene expression in both datasets based on the signed -log(p-value), with the sign depending upon whether a gene was up or down-regulated. We implemented a sliding window approach to scan through genes iteratively in both gene lists and computed hyper-geometric test p-values for each window. The Benjamini-Yekutieli multiple hypothesis correction was applied to the p-values. The computed p-adj values were converted to -log (p-adj) values and represented as a heat map.

## Data and materials availability

Sequencing data will be uploaded to GEO upon acceptance for publication and an accession number will be included in the published version of the article.

## References

1. F. Luo, M. Li, C. Florence, State-Level Economic Costs of Opioid Use Disorder and Fatal Opioid Overdose — United States, 2017. MMWR Morb Mortal Wkly Rep 70, 541–546 (2025).

2. C. Florence, F. Luo, K. Rice, The economic burden of opioid use disorder and fatal opioid overdose in the United States, 2017. Drug Alcohol Depend 218, 108350 (2021).

3. C. Marel, M. Sunderland, K. L. Mills, T. Slade, M. Teesson, C. Chapman, Conditional probabilities of substance use disorders and associated risk factors: Progression from first use to use disorder on alcohol, cannabis, stimulants, sedatives and opioids. Drug Alcohol Depend 194, 136–142 (2019).

4. K. S. Kendler, S. L. Lönn, J. Ektor-Andersen, J. Sundquist, K. Sundquist, Risk factors for the development of opioid use disorder after first opioid prescription: a Swedish national study. Psychol Med 53, 6223–6231 (2023).

5. E. R. Gibney, C. M. Nolan, Epigenetics and gene expression. Heredity (Edinb) 105, 4–13 (2010).

6. C. T. Werner, R. D. Altshuler, Y. Shaham, X. Li, Epigenetic mechanisms in drug relapse. Biol Psychiatry 89, 331 (2021).

7. C. J. Browne, R. Futamura, A. Minier-Toribio, E. M. Hicks, A. Ramakrishnan, F. J. Martínez-Rivera, M. Estill, A. Godino, E. M. Parise, A. Torres-Berrío, A. M. Cunningham, P. J. Hamilton, D. M. Walker, L. M. Huckins, Y. L. Hurd, L. Shen, E. J. Nestler, Transcriptional signatures of heroin intake and relapse throughout the brain reward circuitry in male mice. Sci Adv 9 (2023).

8. S. H. Duttke, P. Montilla-Perez, M. W. Chang, H. Li, H. Chen, L. L. G. Carrette, G. de Guglielmo, O. George, A. A. Palmer, C. Benner, F. Telese, Glucocorticoid Receptor-Regulated Enhancers Play a Central Role in the Gene Regulatory Networks Underlying Drug Addiction. bioRxiv, 2022.01.12.475507 (2022).

9. O. F. Pomerleau, A. C. Collins, S. Shiffman, C. S. Pomerleau, Why some people smoke and others do not: new perspectives. J Consult Clin Psychol 61, 723–731 (1993).

10. O. F. Pomerleau, Individual differences in sensitivity to nicotine: Implications for genetic research on nicotine dependence. Behav Genet 25, 161–177 (1995).

11. J. O’Loughlin, J. DiFranza, R. F. Tyndale, G. Meshefedjian, E. McMillan-Davey, P. B. S. Clarke, J. Hanley, G. Paradis, Nicotine-dependence symptoms are associated with smoking frequency in adolescents. Am J Prev Med 25, 219–225 (2003).

12. J. R. DiFranza, Thwarting science by protecting the received wisdom on tobacco addiction from the scientific method. Harm Reduct J 7, 1–12 (2010).

13. J. R. DiFranza, J. A. Savageau, K. Fletcher, J. O’Loughlin, L. Pbert, J. K. Ockene, A. D. McNeill, J. Hazelton, K. Friedman, G. Dussault, C. Wood, R. J. Wellman, Symptoms of tobacco dependence after brief intermittent use: The development and assessment of nicotine dependence in youth-2 study. Arch Pediatr Adolesc Med 161, 704–710 (2007).

14. M. A. Schuckit, Low level of response to alcohol as a predictor of future alcoholism. American Journal of Psychiatry 151, 184–189 (1994).

15. M. A. Schuckit, T. L. Smith, J. Kalmijn, The search for genes contributing to the low level of response to alcohol: patterns of findings across studies. Alcohol Clin Exp Res 28, 1449–1458 (2004).

16. C. A. Doubeni, G. Reed, J. R. DiFranza, Early Course of Nicotine Dependence in Adolescent Smokers. Pediatrics 125, 1127 (2010).

17. C. J. Morris, S. Mills-Huffnagle, A. E. Zgierska, Can the type of subjective response to first opioid exposure predict the risk of opioid use disorder? A scoping review. Brain Res Bull 188, 67–76 (2022).

18. N. A. Holtz, A. K. Radke, N. E. Zlebnik, A. C. Harris, M. E. Carroll, Intracranial self-stimulation reward thresholds during morphine withdrawal in rats bred for high (HiS) and low (LoS) saccharin intake. Brain Res 1602, 119 (2015).

19. J. M. Wiebelhaus, D. M. Walentiny, P. M. Beardsley, Effects of acute and repeated administration of oxycodone and naloxone-precipitated withdrawal on intracranial self-stimulation in rats. Journal of Pharmacology and Experimental Therapeutics 356, 43–52 (2016).

20. Y. Swain, P. Muelken, A. Skansberg, D. Lanzdorf, Z. Haave, M. G. LeSage, J. C. Gewirtz, A. C. Harris, Higher anhedonia during withdrawal from initial opioid exposure is protective against subsequent opioid self-administration in rats. Psychopharmacology (Berl) 237, 2279 (2020).

21. B. S. Bentzley, K. M. Fender, G. Aston-Jones, The behavioral economics of drug self-administration: a review and new analytical approach for within-session procedures. Psychopharmacology (Berl) 226, 113–125 (2013).

22. W. K. Bickel, M. W. Johnson, M. N. Koffarnus, J. MacKillop, J. G. Murphy, The Behavioral Economics of Substance Use Disorders: reinforcement pathologies and their repair. Annu Rev Clin Psychol 10, 641 (2014).

23. I. Fredriksson, M. Venniro, D. J. Reiner, J. J. Chow, J. M. Bossert, Y. Shaham, Animal Models of Drug Relapse and Craving after Voluntary Abstinence: A Review. Pharmacol Rev 73, 1050–1083 (2021).

24. L. Chen, T. Kreko-Pierce, S. L. Cassoday, L. Al-Harthi, X. T. Hu, Methamphetamine self-administration causes neuronal dysfunction in rat medial prefrontal cortex in a sex-specific and withdrawal time-dependent manner. Front Pharmacol 16, 1527795 (2025).

25. M. Yu, L. Lin, K. Xu, M. Xu, J. Ren, X. Niu, X. Gao, M. Zhang, Z. Yang, J. Dang, Q. Tao, S. Han, W. Wang, J. Cheng, Y. Zhang, Changes in aspartate metabolism in the medial-prefrontal cortex of nicotine addicts based on J-edited magnetic resonance spectroscopy. Hum Brain Mapp 44, 6429–6438 (2023).

26. X. Yang, Y. J. Meng, Y. J. Tao, R. H. Deng, H. Y. Wang, X. J. Li, W. Wei, Y. Hua, Q. Wang, W. Deng, L. S. Zhao, X. H. Ma, M. L. Li, J. J. Xu, J. Li, Y. S. Liu, Z. Tang, X. D. Du, J. W. Coid, A. J. Greenshaw, T. Li, W. J. Guo, Functional Connectivity of Nucleus Accumbens and Medial Prefrontal Cortex With Other Brain Regions During Early-Abstinence Is Associated With Alcohol Dependence and Relapse: A Resting-Functional Magnetic Resonance Imaging Study. Front Psychiatry 12, 609458 (2021).

27. T. G. Brown, J. Xu, Y. L. Hurd, Y. X. Pan, Dysregulated expression of the alternatively spliced variant mRNAs of the mu opioid receptor gene, OPRM1, in the medial prefrontal cortex of male human heroin abusers and heroin self-administering male rats. J Neurosci Res 100, 35–47 (2022).

28. S. K. Abdulaev, D. A. Tarumov, V. K. Shamrey, A. G. Trufanov, N. A. Puchkov, K. V. Markin, Y. E. Prochik, Functional Impairments in the Large-Scale Resting Networks of the Brain in Opioid Addiction. Neurosci Behav Physiol 53, 1502–1508 (2023).

29. A. C. Harris, P. Muelken, J. R. Smethells, M. Krueger, L. S. Le, Similar precipitated withdrawal effects on intracranial self-stimulation during chronic infusion of an e-cigarette liquid or nicotine alone. Pharmacol Biochem Behav 161, 1–5 (2017).

30. S. Vlachou, A. Markou, Intracranial self-stimulation. Neuromethods 53, 3–56 (2011).

31. S. R. Hursh, A. Silberberg, Economic Demand and Essential Value. Psychol Rev 115, 186–198 (2008).

32. D. M. Walker, H. M. Cates, Y. H. E. Loh, I. Purushothaman, A. Ramakrishnan, K. M. Cahill, C. K. Lardner, A. Godino, H. G. Kronman, J. Rabkin, Z. S. Lorsch, P. Mews, M. A. Doyle, J. Feng, B. Labonté, J. W. Koo, R. C. Bagot, R. W. Logan, M. L. Seney, E. S. Calipari, L. Shen, E. J. Nestler, Cocaine Self-administration Alters Transcriptome-wide Responses in the Brain’s Reward Circuitry. Biol Psychiatry 84, 867–880 (2018).

33. S. X. Liu, M. S. Gades, Y. Swain, A. Ramakrishnan, A. C. Harris, P. V. Tran, J. C. Gewirtz, Repeated morphine exposure activates synaptogenesis and other neuroplasticity-related gene networks in the dorsomedial prefrontal cortex of male and female rats. Drug Alcohol Depend 221, 108598 (2021).

34. G. E. Hoffman, E. E. Schadt, variancePartition: Interpreting drivers of variation in complex gene expression studies. BMC Bioinformatics 17, 1–13 (2016).

35. T. Fuller, P. Langfelder, A. Presson, S. Horvath, Review of Weighted Gene Coexpression Network Analysis. Handbook of Statistical Bioinformatics, 369–388 (2011).

36. P. Langfelder, R. Luo, M. C. Oldham, S. Horvath, Is My Network Module Preserved and Reproducible? PLoS Comput Biol 7, e1001057 (2011).

37. A. A. Margolin, I. Nemenman, K. Basso, C. Wiggins, G. Stolovitzky, R. D. Favera, A. Califano, ARACNE: An algorithm for the reconstruction of gene regulatory networks in a mammalian cellular context. BMC Bioinformatics 7, 1–15 (2006).

38. K. Scott, R. O’Rourke, A. Gillen, B. Appel, Prdm8 regulates pMN progenitor specification for motor neuron and oligodendrocyte fates by modulating the Shh signaling response. Development 147 (2020).

39. H. Fu, S. Kesari, J. Cai, Tcf7l2 is Tightly Controlled During Myelin Formation. Cell Mol Neurobiol 32, 345 (2012).

40. S. Zhang, X. Zhu, X. Gui, C. Croteau, L. Song, J. Xu, A. Wang, P. Bannerman, F. Guo, Sox2 Is Essential for Oligodendroglial Proliferation and Differentiation during Postnatal Brain Myelination and CNS Remyelination. The Journal of Neuroscience 38, 1802 (2018).

41. A. Benraiss, J. N. Mariani, A. Tate, P. M. Madsen, K. M. Clark, K. A. Welle, R. Solly, L. Capellano, K. Bentley, D. Chandler-Militello, S. A. Goldman, A TCF7L2-responsive suppression of both homeostatic and compensatory remyelination in Huntington disease mice. Cell Rep 40 (2022).

42. B. N. Kuhn, N. Cannella, A. S. Chitre, K. M. H. Nguyen, K. Cohen, D. Chen, B. Peng, K. S. Ziegler, B. Lin, B. B. Johnson, T. Missfeldt Sanches, A. D. Crow, V. Lunerti, A. Gupta, E. Dereschewitz, L. Soverchia, J. L. Hopkins, A. T. Roberts, M. Ubaldi, S. Abdulmalek, A. Kinen, G. Hardiman, D. Chung, O. Polesskaya, L. C. Solberg Woods, R. Ciccocioppo, P. W. Kalivas, A. A. Palmer, Genome-wide association study reveals multiple loci for nociception and opioid consumption behaviors associated with heroin vulnerability in outbred rats. Mol Psychiatry 30, 3363–3375 (2025).

43. M. L. Seney, S. M. Kim, J. R. Glausier, M. A. Hildebrand, X. Xue, W. Zong, J. Wang, M. A. Shelton, B. D. N. Phan, C. Srinivasan, A. R. Pfenning, G. C. Tseng, D. A. Lewis, Z. Freyberg, R. W. Logan, Transcriptional Alterations in Dorsolateral Prefrontal Cortex and Nucleus Accumbens Implicate Neuroinflammation and Synaptic Remodeling in Opioid Use Disorder. Biol Psychiatry 90, 550–562 (2021).

44. M. Hearing, Prefrontal-accumbens opioid plasticity: Implications for relapse and dependence. Academic Press [Preprint] (2019). 10.1016/j.phrs.2018.11.012.

45. V. Pascoli, J. Terrier, J. Espallergues, E. Valjent, E. C. O’connor, C. Lüscher, Contrasting forms of cocaine-evoked plasticity control components of relapse. Nature 509, 459–464 (2014).

46. É. Kestering-Ferreira, B. A. Heberle, F. Sindermann Lumertz, P. H. Gobira, R. Orso, R. Grassi-Oliveira, T. W. Viola, Sex differences in sensitivity to fentanyl effects in mice: Behavioral and molecular findings during late adolescence. Neurosci Lett 837, 137898 (2024).

47. G. Picci, D. H. Fishbein, J. W. VanMeter, E. J. Rose, Effects of OPRM1 and DRD2 on brain structure in drug-naïve adolescents: Genetic and neural vulnerabilities to substance use. Psychopharmacology (Berl) 239, 141–152 (2022).

48. D. Nishizawa, K. Fukuda, S. Kasai, J. Hasegawa, Y. Aoki, A. Nishi, N. Saita, Y. Koukita, M. Nagashima, R. Katoh, Y. Satoh, M. Tagami, S. Higuchi, H. Ujike, N. Ozaki, T. Inada, N. Iwata, I. Sora, M. Iyo, N. Kondo, M. J. Won, N. Naruse, K. Uehara-Aoyama, M. Itokawa, M. Koga, T. Arinami, Y. Kaneko, M. Hayashida, K. Ikeda, Genome-wide association study identifies a potent locus associated with human opioid sensitivity. Molecular Psychiatry 2014 19:1 19, 55–62 (2012).

49. K. Ikeda, S. Ide, W. Han, M. Hayashida, G. R. Uhl, I. Sora, How individual sensitivity to opiates can be predicted by gene analyses. Trends Pharmacol Sci 26, 311–317 (2005).

50. K. D. Ersche, C. Meng, H. Ziauddeen, J. Stochl, G. B. Williams, E. T. Bullmore, T. W. Robbins, Brain networks underlying vulnerability and resilience to drug addiction. Proc Natl Acad Sci U S A 117, 15253–15261 (2020).

51. J. Cuitavi, J. V. Torres-Pérez, J. D. Lorente, Y. Campos-Jurado, P. Andrés-Herrera, A. Polache, C. Agustín-Pavón, L. Hipólito, Crosstalk between Mu-Opioid receptors and neuroinflammation: Consequences for drug addiction and pain. Neurosci Biobehav Rev 145, 105011 (2023).

52. C. M. Cahill, A. M. Taylor, Neuroinflammation—a co-occurring phenomenon linking chronic pain and opioid dependence. Curr Opin Behav Sci 13, 171–177 (2017).

53. M. Kohno, J. Link, L. E. Dennis, H. McCready, M. Huckans, W. F. Hoffman, J. M. Loftis, Neuroinflammation in addiction: A review of neuroimaging studies and potential immunotherapies. Pharmacol Biochem Behav 179, 34–42 (2019).

54. C. A. McClung, E. J. Nestler, Regulation of gene expression and cocaine reward by CREB and ΔFosB. Nat Neurosci 6, 1208–1215 (2003).

55. E. J. Nestler, Molecular neurobiology of addiction. American Journal on Addictions 10, 201–217 (2001).

56. Y. Yang, L. Guo, L. Chen, B. Gong, D. Jia, Q. Sun, Nuclear transport proteins: structure, function and disease relevance. Signal Transduction and Targeted Therapy 2023 8:1 8, 1–29 (2023).

57. M. A. D’angelo, Nuclear pore complexes as hubs for gene regulation. Nucleus 9, 142–148 (2018).

58. L. Chen, Y. Wang, J. Lin, Z. Song, Q. Wang, W. Zhao, Y. Wang, X. Xiu, Y. Deng, X. Li, Q. Li, X. Wang, J. Li, X. Liu, K. Liu, J. Zhou, K. Li, Y. Liu, S. Liao, Q. Deng, C. Xu, Q. Sun, S. Wu, K. Zhang, M. X. Guan, T. Zhou, F. Sun, X. Cai, C. Huang, G. Shan, Exportin 4 depletion leads to nuclear accumulation of a subset of circular RNAs. Nature Communications 2022 13:1 13, 1–18 (2022).

59. D. A. Jans, A. J. Martin, K. M. Wagstaff, Inhibitors of nuclear transport. Curr Opin Cell Biol 58, 50–60 (2019).

60. Z. Boudhraa, E. Carmona, D. Provencher, A. M. Mes-Masson, Ran GTPase: A Key Player in Tumor Progression and Metastasis. Front Cell Dev Biol 8 (2020).

61. Q. Ru, Y. Wang, E. Zhou, L. Chen, Y. Wu, The potential therapeutic roles of Rho GTPases in substance dependence. Front Mol Neurosci 16, 1125277 (2023).

62. W. Wang, Y. Y. Ju, Q. X. Zhou, J. X. Tang, M. Li, L. Zhang, S. Kang, Z. G. Chen, Y. J. Wang, H. Ji, Y. Q. Ding, L. Xu, J. G. Liu, The small GTPase rac1 contributes to extinction of aversive memories of drug withdrawal by facilitating GABAa receptor endocytosis in the vmPFC. Journal of Neuroscience 37, 7096–7110 (2017).

63. M. H. S. Deurloo, S. Eide, E. Turlova, Q. Li, S. Spijker, H. S. Sun, A. J. A. Groffen, Z. P. Feng, Rasal1 regulates calcium dependent neuronal maturation by modifying microtubule dynamics. Cell Biosci 14 (2024).

64. F. Di Tullio, M. Schwarz, H. Zorgati, S. Mzoughi, E. Guccione, The duality of PRDM proteins: epigenetic and structural perspectives. FEBS J 289, 1256–1275 (2022).

65. S. Marques, A. Zeisel, S. Codeluppi, D. Van Bruggen, A. M. Falcão, L. Xiao, H. Li, M. Häring, H. Hochgerner, R. A. Romanov, D. Gyllborg, A. B. Muñoz-Manchado, G. La Manno, P. Lönnerberg, E. M. Floriddia, F. Rezayee, P. Ernfors, E. Arenas, J. Hjerling-Leffler, T. Harkany, W. D. Richardson, S. Linnarsson, G. Castelo-Branco, Oligodendrocyte heterogeneity in the mouse juvenile and adult central nervous system. Science (1979) 352, 1326–1329 (2016).

66. C. Zhao, D. Ma, M. Zawadzka, S. P. J. Fancy, L. Elis-Williams, G. Bouvier, J. H. Stockley, G. M. de Castro, B. Wang, S. Jacobs, P. Casaccia, R. J. M. Franklin, Sox2 sustains recruitment of oligodendrocyte progenitor cells following CNS demyelination and primes them for differentiation during remyelination. Journal of Neuroscience 35, 11482–11499 (2015).

67. A. Kruyer, A. Angelis, C. Garcia-Keller, H. Li, P. W. Kalivas, Plasticity in astrocyte subpopulations regulates heroin relapse. Sci Adv 8, 7044 (2022).

68. A. Kruyer, P. W. Kalivas, Astrocytes as cellular mediators of cue reactivity in addiction. Curr Opin Pharmacol 56, 1–6 (2021).

69. A. Kruyer, P. W. Kalivas, M. D. Scofield, Astrocyte regulation of synaptic signaling in psychiatric disorders. Neuropsychopharmacology 2022 48:1 48, 21–36 (2022).

70. B. Yalçın, M. B. Pomrenze, K. Malacon, R. Drexler, A. E. Rogers, K. Shamardani, I. J. Chau, K. R. Taylor, L. Ni, D. Contreras-Esquivel, R. C. Malenka, M. Monje, Myelin plasticity in the ventral tegmental area is required for opioid reward. Nature 2024 630:8017 630, 677–685 (2024).

71. R. A. Hill, A. Nishiyama, E. G. Hughes, Features, Fates, and Functions of Oligodendrocyte Precursor Cells. Cold Spring Harb Perspect Biol 16 (2024).

72. S. O. Spitzer, S. Sitnikov, Y. Kamen, K. A. Evans, D. Kronenberg-Versteeg, S. Dietmann, O. de Faria, S. Agathou, R. T. Káradóttir, Oligodendrocyte Progenitor Cells Become Regionally Diverse and Heterogeneous with Age. Neuron 101, 459–471.e5 (2019).

73. W. Xin, M. Kaneko, R. H. Roth, A. Zhang, S. Nocera, J. B. Ding, M. P. Stryker, J. R. Chan, Oligodendrocytes and myelin limit neuronal plasticity in visual cortex. Nature 2024 633:8031 633, 856–863 (2024).

74. J. Simino, Y. J. Sung, R. Kume, K. Schwander, D. C. Rao, Gene-alcohol interactions identify several novel blood pressure loci including a promising locus near SLC16A9. Front Genet 4, 277 (2013).

75. M. Argos, L. Tong, B. L. Pierce, M. Rakibuz-Zaman, A. Ahmed, T. Islam, M. Rahman, R. Paul-Brutus, R. Rahaman, S. Roy, F. Jasmine, M. G. Kibriya, H. Ahsan, Genome-wide association study of smoking behaviors among Bangladeshi adults. J Med Genet 51, 327 (2014).

76. M. Zaupa, N. Nagaraj, A. Sylenko, H. Baier, S. Sawamiphak, A. Filosa, The Calmodulin-interacting peptide Pcp4a regulates feeding state-dependent behavioral choice in zebrafish. Neuron 112, 1150–1164.e6 (2024).

77. S. Kourrich, P. E. Rothwell, J. R. Klug, M. J. Thomas, Cocaine experience controls bidirectional synaptic plasticity in the nucleus accumbens. Journal of Neuroscience 27, 7921–7928 (2007).

78. H. M. Cates, X. Li, I. Purushothaman, P. J. Kennedy, L. Shen, Y. Shaham, E. J. Nestler, Genome-wide transcriptional profiling of central amygdala and orbitofrontal cortex during incubation of methamphetamine craving. Neuropsychopharmacology 43, 2426 (2018).

79. H. L. Mayberry, C. C. Bavley, R. Karbalaei, D. R. Peterson, A. R. Bongiovanni, A. S. Ellis, S. H. Downey, A. B. Toussaint, M. E. Wimmer, Transcriptomics in the nucleus accumbens shell reveal sex- and reinforcer-specific signatures associated with morphine and sucrose craving. Neuropsychopharmacology 47, 1764–1775 (2022).

80. M. A. Shelton, N. Horan, X. Xue, L. Maturin, D. Eacret, J. Michaud, N. Singh, B. R. Williams, M. C. Gamble, J. A. Seggio, M. K. Fish, B. D. N. Phan, G. C. Tseng, J. A. Blendy, L. C. Solberg Woods, A. A. Palmer, O. George, R. W. Logan, M. L. Seney, Sex-Specific Concordance of Striatal Transcriptional Signatures of Opioid Addiction in Human and Rodent Brains. Biological Psychiatry Global Open Science 5, 100476 (2025).

81. N. A. Holtz, A. K. Radke, N. E. Zlebnik, A. C. Harris, M. E. Carroll, Intracranial self-stimulation reward thresholds during morphine withdrawal in rats bred for high (HiS) and low (LoS) saccharin intake. Brain Res 1602, 119 (2015).

82. J. Liu, G. Schulteis, Brain reward deficits accompany naloxone-precipitated withdrawal from acute opioid dependence. Pharmacol Biochem Behav 79, 101–108 (2004).

83. Y. Swain, P. Muelken, A. Skansberg, D. Lanzdorf, Z. Haave, M. G. LeSage, J. C. Gewirtz, A. C. Harris, Higher anhedonia during withdrawal from initial opioid exposure is protective against subsequent opioid self-administration in rats. Psychopharmacology (Berl) 237, 2279 (2020).

84. J. Feng, M. Wilkinson, X. Liu, I. Purushothaman, D. Ferguson, V. Vialou, I. Maze, N. Shao, P. Kennedy, J. W. Koo, C. Dias, B. Laitman, V. Stockman, Q. LaPlant, M. E. Cahill, E. J. Nestler, L. Shen, Chronic cocaine-regulated epigenomic changes in mouse nucleus accumbens. Genome Biol 15 (2014).

85. G. E. Hodes, M. L. Pfau, I. Purushothaman, H. F. Ahn, S. A. Golden, D. J. Christoffel, J. Magida, A. Brancato, A. Takahashi, M. E. Flanigan, C. Menard, H. Aleyasin, J. W. Koo, Z. S. Lorsch, J. Feng, M. Heshmati, M. Wang, G. Turecki, R. Neve, B. Zhang, L. Shen, E. J. Nestler, S. J. Russo, Sex Differences in Nucleus Accumbens Transcriptome Profiles Associated with Susceptibility versus Resilience to Subchronic Variable Stress. Journal of Neuroscience 35, 16362–16376 (2015).

86. M. Li, P. Xu, Y. Xu, H. Teng, W. Tian, Q. Du, M. Zhao, Dynamic expression changes in the transcriptome of the prefrontal cortex after repeated exposure to cocaine in mice. Front Pharmacol 8, 142 (2017).

87. D. M. Walker, H. M. Cates, Y. H. E. Loh, I. Purushothaman, A. Ramakrishnan, K. M. Cahill, C. K. Lardner, A. Godino, H. G. Kronman, J. Rabkin, Z. S. Lorsch, P. Mews, M. A. Doyle, J. Feng, B. Labonté, J. W. Koo, R. C. Bagot, R. W. Logan, M. L. Seney, E. S. Calipari, L. Shen, E. J. Nestler, Cocaine Self-administration Alters Transcriptome-wide Responses in the Brain’s Reward Circuitry. Biol Psychiatry 84, 867–880 (2018).

88. Y. Swain, P. Muelken, M. G. LeSage, J. C. Gewirtz, A. C. Harris, Locomotor activity does not predict individual differences in morphine self-administration in rats. Pharmacol Biochem Behav 166, 48 (2018).

89. D. Kim, J. M. Paggi, C. Park, C. Bennett, S. L. Salzberg, Graph-Based Genome Alignment and Genotyping with HISAT2 and HISAT-genotype. Nat Biotechnol 37, 907 (2019).

90. M. I. Love, W. Huber, S. Anders, Moderated estimation of fold change and dispersion for RNA-seq data with DESeq2. Genome Biol 15 (2014).

91. A. Barks, S. J. B. Fretham, M. K. Georgieff, P. V. Tran, Early-Life Neuronal-Specific Iron Deficiency Alters the Adult Mouse Hippocampal Transcriptome. Journal of Nutrition 148, 1521–1528 (2018).

92. S. B. Plaisier, R. Taschereau, J. A. Wong, T. G. Graeber, Rank–rank hypergeometric overlap: identification of statistically significant overlap between gene-expression signatures. Nucleic Acids Res 38, e169 (2010).

93. A. C. Harris, P. Muelken, S. X. Liu, J. R. Smethells, M. G. LeSage, J. C. Gewirtz, Magnitude and predictors of elasticity of demand for morphine are similar in male and female rats. Front Behav Neurosci 18, 1443364 (2024).

