## Supplementary Materials for "Transcriptional signatures in the rat medial prefrontal cortex associated with vulnerability and resilience across distinct phases of opioid use disorder"

Science Advances
AAAS

### Supplementary Materials for

#### Transcriptional signatures in the rat medial prefrontal cortex associated with vulnerability and resilience across distinct phases of opioid use disorder

Shirelle X. Liu *et al.*

**This PDF file includes:**

Supplementary Text
Figs. S1 to S6
Tables S1 to S6
References (1 to 5)

#### Supplementary Text

#### Supplementary Methods

##### *Apparatus*

Intracranial self-stimulation (ICSS). Rats were tested in operant conditioning chambers (29×26×33 cm; Med Associates, St. Albans, VT, USA) placed inside sound-attenuating cubicles. A 5-cm-wide metal wheel manipulandum was fixed to the front wall. Brain stimulation was administered with constant current stimulators (model #PHM-152, Med Associates). Rats were connected to the stimulation circuit through bipolar leads (Plastics One, Roanoke, VA, USA) attached to gold-contact swivel commutators (Plastics One). MED-PC IV software was used to control stimulation parameters and for data collection.

Morphine self-administration (Morphine SA). Morphine SA sessions were conducted using standard operant conditioning chambers (model ENV-007, Med Associates, Inc). Each chamber contained two response levers, a green light emitting diode (LED) or white stimulus (i.e., cue) light located 2 cm above each lever, and a house light that provided ambient illumination. Each chamber was placed inside a sound-attenuating cubicle equipped with an exhaust fan that provided masking noise. An infusion pump (model PHM-100-15, Med Associates) placed outside each cubicle delivered infusions in a volume of 0.1 ml/kg over approximately 1 second. MED-PC IV software (Med Associates) was used for operating the experimental apparatus and recording data.

##### *Surgery*

ICSS. Animals were anesthetized using inhaled isoflurane (1-3%) and implanted with a bipolar stainless-steel electrode (Plastics One) in the medial forebrain bundle at the level of the lateral hypothalamus using our standard procedures (e.g., Roiko et al., 2009; Harris et al., 2010) (*1*, *2*). Animals were allowed to recover for at least 1 week prior to ICSS training. During the first 2 days of recovery, all animals received injections of the antibiotic ceftriaxone (5.25 mg, i.m.) and the analgesic buprenorphine (0.1 mg/kg, s.c.).

Morphine SA. Each rat was implanted with a chronic indwelling catheter into the right jugular vein under isoflurane (1-3%) anesthesia, using general surgical procedures described in detail elsewhere (*3*, *4*). To allow for morphine SA, the catheter was either externalized between the scapulae and attached to a vascular-access harness (VAH95AB, Instech Laboratories, Plymouth Meeting, PA) or attached to an indwelling vascular-access button (VABR2B, Instech Laboratories). In either case, the catheter was connected mid-scapulae to a tether that ran through a fluid swivel before connecting to the drug pump. Animals were allowed to recover for one week after surgery, during which time they received daily i.v. infusions of heparinized saline, ceftriaxone antibiotic (5.25 mg, first three days only), and s.c. injections of buprenorphine (0.05 mg/kg; first two days only) for analgesia. Infusions of methohexital (0.1 ml, 10 mg/ml, i.v.) were administered to check catheter patency post-session on Fridays. If a catheter became occluded (indicated by a failure of the animal to exhibit anesthesia within 3-5 sec after methohexital infusion), another catheter was implanted into the ipsilateral femoral vein. If that catheter failed, a third catheter was implanted into the contralateral femoral vein. Failure of the third catheter resulted in removal of the animal from the study.

##### *General testing procedures*

ICSS. Each trial was initiated with presentation of a non-contingent stimulus (0.1-ms cathodal square wave pulses at a frequency of 100 Hz for 500 ms) followed by a 7.5-s window, during which a positive response on the wheel manipulandum produced a second contingent stimulation

identical to the first. Lack of responding during the 7.5-s window was considered a negative response. Each positive or negative response was followed by a variable inter-trial interval averaging 10 s (range, 7.5–12.5 s), during which time additional responses delayed the onset of the subsequent trial by 12.5 s. Stimulus intensities were presented in four alternating descending and ascending series (step size, 5 μA), with five trials presented at each current intensity step. The current threshold for each series was defined as the midpoint between two consecutive intensity steps that yielded three or more positive responses and two consecutive intensity steps that yielded three or more negative responses. The overall ICSS threshold for the session was defined as the mean of the current thresholds from the four alternating series. To assess performance effects (e.g., motor disruption), response latencies (time between onset of the non-contingent stimulus and a positive response) were averaged across all trials in which a positive response was made.
Morphine SA. During each 2-hr session during, responding on the "active" response lever resulted in an i.v. infusion of morphine sulfate (0.4 mg/kg/inf) that was accompanied by offset of the cue light above the active response lever. Following a 5-second timeout period, the cue light above the active lever was illuminated to signal availability of the next infusion. Responses on the other response lever (the "inactive" lever) were recorded but had no programmed consequences. This unit dose and access duration support reliable morphine SA in the absence of self-mutilation associated with higher unit doses and longer sessions (*5*). On the first day of morphine SA, food powder was placed on the active lever to facilitate contact with the lever. Data from this session were not included in the data analysis.

**Supplementary Statistics**

Withdrawal-induced anhedonia. Baseline ICSS thresholds (in μA) and response latencies (in sec) were defined as the mean during the last five sessions prior to withdrawal testing. These data were analyzed using separate 2-factor ANOVAs with morphine dose (0 or 5.6 mg/kg) and sex as between-subject factors. ICSS response latencies during withdrawal testing were subsequently expressed as a percentage of baseline and analyzed using a 3-factor ANOVA with morphine dose and sex as between-subject factors and session as a within-subject factor.

Demand. Infusions per day during the last 5 sessions of acquisition were compared using 3-factor ANOVA with morphine unit dose (0 or 0.4 mg/kg/infusion) and sex as between-subject factors and session as a within-subject factor.

Reinstatement. To assess differences in baseline SA levels of saline vs morphine, a three-factor ANOVA (sex x morphine dose x acquisition day) was conducted on active lever presses across the final five sessions prior to meeting acquisition criteria. The same analysis was conducted during the first 7 days of extinction to determine if responding initially underwent extinction in either drug group (saline vs morphine). Finally, to confirm that a significant reduction in responding occurred during extinction in the morphine SA group, but not the saline group, active lever responding was averaged across the final five sessions of acquisition and extinction for each group analyzed using a two-factor ANOVA within phase (acquisition versus extinction) and between SA group (saline vs morphine).

#### Supplementary Results

***Experiment 1: Baseline ICSS measures***

There was no effect of morphine dose, sex, or interaction between these factors for either baseline ICSS thresholds or baseline response latencies (Table S1).

***Experiment 1: ICSS response latencies***

There was a significant effect of session on ICSS response latencies $F(5.1, 180.5) = 2.3$, $p < 0.05$), but no effect of morphine dose, sex, or interaction between these variables (Figure S1A).

***Experiment 2: Acquisition of MSA.***

A 3-factor ANOVA on infusions per session during the final 5 sessions of acquisition indicated a significant main effect of morphine dose $F(1, 30) = 19.6$, $p < 0.0001$), but no significant effects of sex, session, or interaction between these variables (Fig S1B).

**Experiment 3: Acquisition and extinction of MSA prior to reinstatement.**

Analysis of baseline levels of SA indicated a significant main effect of morphine dose $F(1,88) = 21.24$, $p < 0.0001$ and an interaction of sex and morphine dose $F(1,88) = 6.47$, $p < 0.05$. Post-hoc tests found significantly higher responding in the Morphine group vs saline and that males responded significantly more in the saline control group than females. A 3-factor ANOVA of active lever responding during the first week of extinction , revealed a main effect of morphine dose $F(1, 88) = 4.736$, $p < 0.05$, session $F(3.63, 319) = 7.33$, $p < 0.0001$ and an interaction of these two factors $F(6, 528) = 4.30$, $p < 0.0001$, but no main effects of sex or interaction with sex. After collapsing across sex, a 2-factor ANOVA follow-up found a main effect of morphine dose $F(1, 90) = 4.655$, $p < .05$, Session $F(3.56, 320.2) = 7.152$, $p < 0.0001$ and the interaction of these factors $F(6, 540) = 4.197$, $p < 0.001$ and that this occurred primarily during the first session according to a Šídákpost-hoc test. Active lever responding was also significantly lower following 21 sessions of extinction. A comparison of mean active responding across the final five sessions of the acquisition and extinction phases using a two-way ANOVA (Phase x SA drug) revealed a significant effect of SA drug $F(1, 90) = 4.71$, $p < 0.05$ and an interaction of Phase x SA drug $F(1, 90) = 21.41$, $p < 0.0001$. Post-hoc tests showed a significant reduction in responding from the acquisition baseline in the morphine SA group, but not the saline SA group.

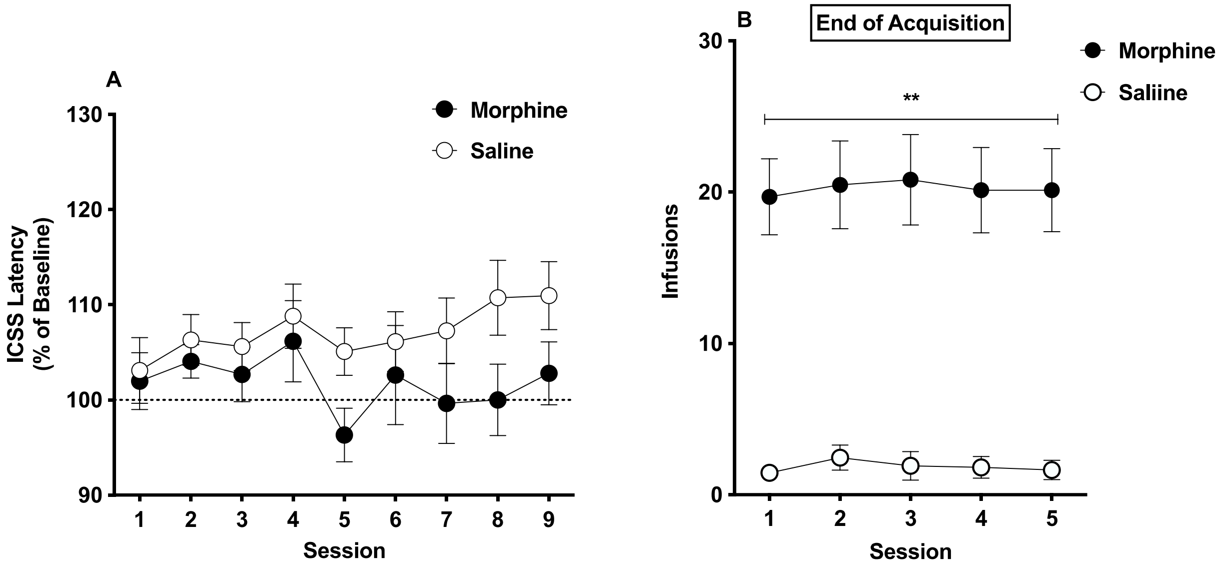


**Fig. S1.**

(A) ICSS response latencies (% Baseline, Mean ± SEM) during spontaneous morphine withdrawal in Experiment 1. (B) Mean (± SEM) infusions per session (Mean ± SEM) during the final 5 sessions of acquisition in Experiment 2. ** Different from Saline, $p < 0.0001$.

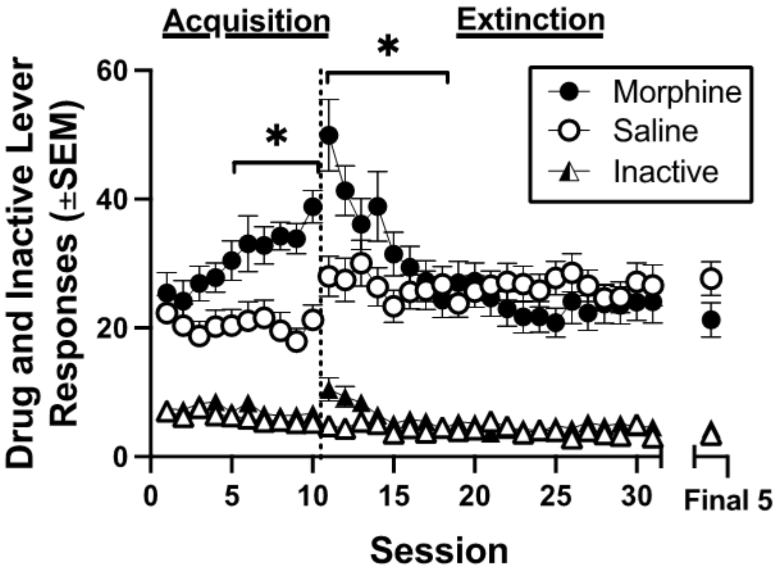


**Fig. S2.**

Shows mean (± SEM) active (drug associated; circles) lever and inactive lever responses (triangles) across the last 10 sessions of acquisition and across the first 21 sessions of extinction in rats responding for saline (open symbols) and 0.4 mg/kg morphine (filled symbols). "Final 5" represents the average responding across the final 5 sessions of extinction (the length of extinction assessment was from 21 to 42 until extinction criteria were met). * Indicates a significant difference from saline controls.

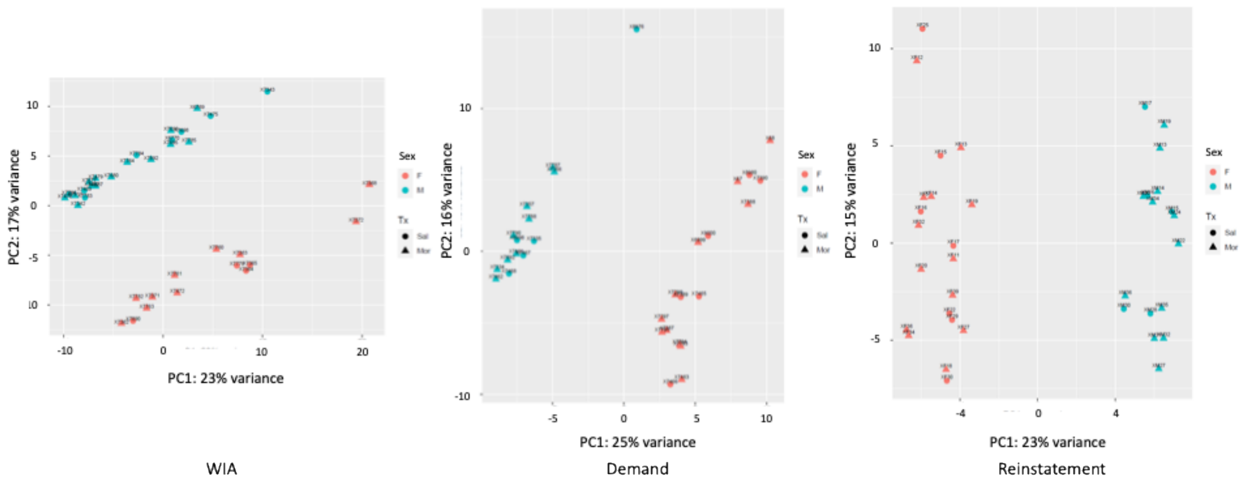


**Fig. S3.**
PCA plots with sex and treatment (Tx) as two dimensions, showing a clear sex separation in three paradigms.

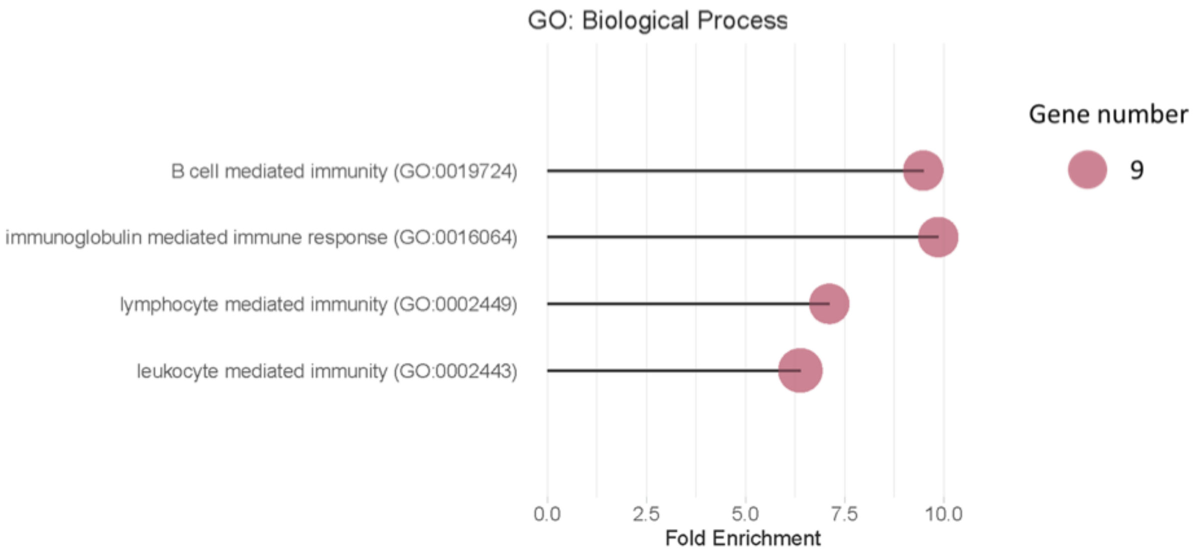


**Fig. S4.**

A WGCNA "White" module with decreased connectivity in Morphine rats showed enrichment of genes involved in neuroimmune function.

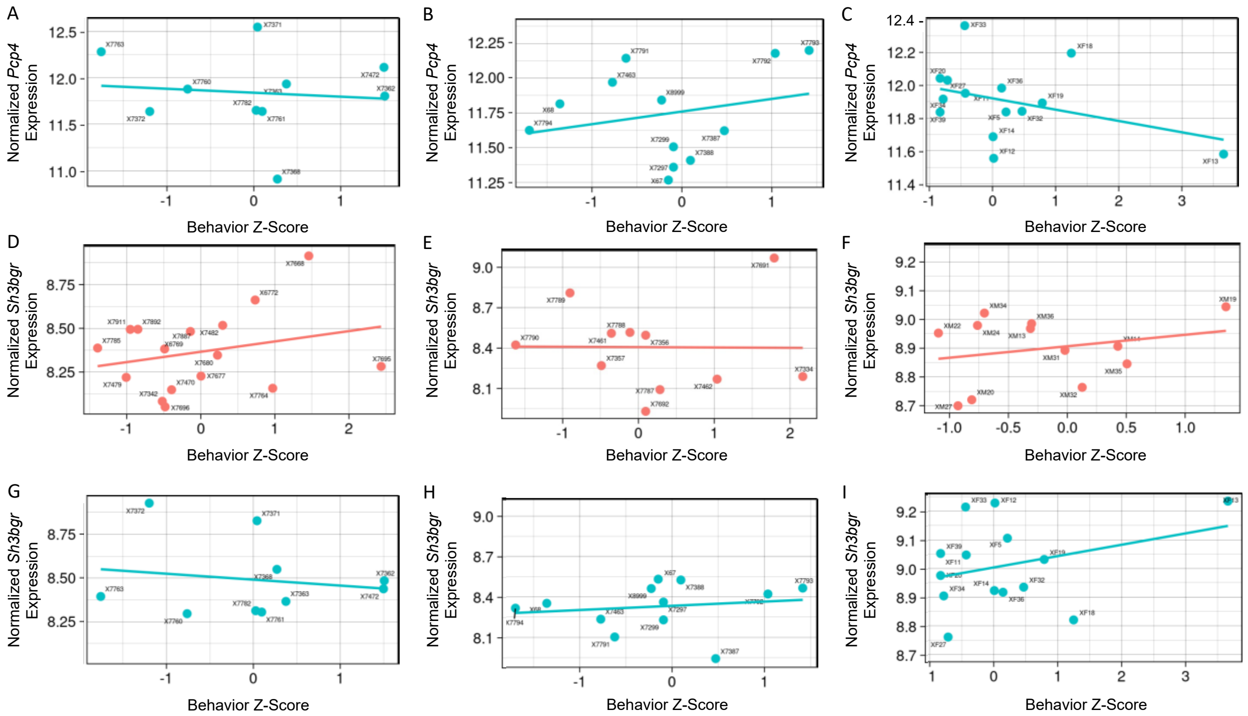

**Fig. S5.**

Coherence with other rat datasets. (A-C) Correlation between normalized *Pcp4* expression and behavioral Z-Scores in female WIA (A), Demand (B), and Reinstatement (C) rats. (D-F) Correlation between normalized *Sh3bgr* expression and behavioral Z-Scores in male WIA (D), Demand (E), and Reinstatement (F) rats. (G-I) Correlation between normalized *Sh3bgr* expression and behavioral Z-Scores in female WIA (G), Demand (H), and Reinstatement (I) rats.

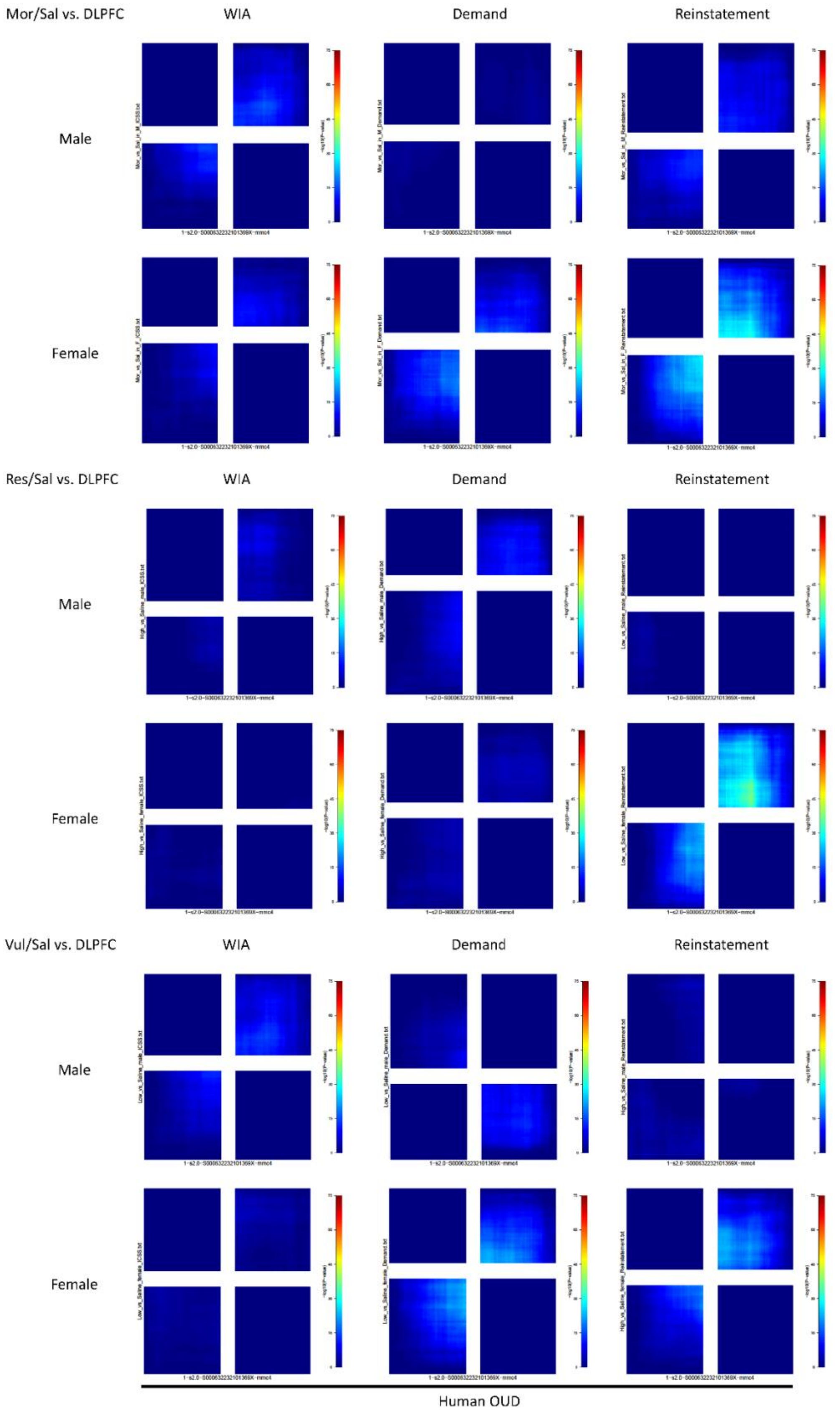
Mor/Sal vs. DLPFC
WIA
Demand
Reinstatement
Male
Female
Res/Sal vs. DLPFC
WIA
Demand
Reinstatement
Male
Female
Vul/Sal vs. DLPFC
WIA
Demand
Reinstatement
Male
Female
Human OUD

**Fig. S6.**

RRHO comparing differential gene expression in the dlPFC of the human OUD patients vs. controls and morphine-exposed, resilient, or vulnerable vs. control rats in in the three paradigms.

| | Male | | Female | |
|---|---|---|---|---|
| | Threshold (µA) | Latency (sec) | Threshold (µA) | Latency (sec) |
| Morphine | 78.1 ± 8.8 | 2.6 ± 0.1 | 88.6 ± 10.1 | 2.7 ± 0.2 |
| Saline | 87.2 ± 12.6 | 2.7 ± 0.1 | 75.1 ± 7.9 | 2.4 ± 0.2 |

**Table S1.**

Mean (±SEM) baseline ICSS thresholds (in µA) and response latencies (in sec) in males and females administered morphine or saline in Experiment 1.

| Subject | α | $Q_0$ | $R^2$ |
|---|---|---|---|
| **Males** | | | |
| 1 | 0.0012 | 10.0 | 0.92 |
| 2 | 0.00099 | 7.3 | 0.96 |
| 3 | 0.00057 | 7.6 | 0.96 |
| 4 | 0.00019 | 33.0 | 0.84 |
| 5 | 0.00086 | 6.9 | 0.95 |
| 6 | 0.0016 | 4.8 | 0.90 |
| 7 | 0.002 | 12.0 | 0.92 |
| 8 | 0.0011 | 8.3 | 0.98 |
| 9 | 0.0022 | 5.8 | 0.94 |
| 10 | 0.0011 | 9.1 | 0.92 |
| 11 | 0.00079 | 8.7 | 0.95 |
| **Mean** | **0.00115** | **10.31** | **0.93** |
| **SEM** | **0.00018** | **2.34** | **0.01** |
| **Females** | | | |
| 1 | 0.00072 | 8.3 | 0.94 |
| 2 | 0.0016 | 9.4 | 0.98 |
| 3 | 0.0018 | 12.0 | 0.90 |
| 4 | 0.00015 | 20.0 | 0.94 |
| 5 | 0.00064 | 17.0 | 0.89 |
| 6 | 0.0013 | 5.6 | 0.93 |
| 7 | 0.0011 | 8.8 | 0.94 |
| 8 | 0.001 | 10.0 | 0.97 |
| 9 | 0.001 | 10.0 | 0.95 |
| 10 | 0.00093 | 8.6 | 0.88 |
| 11 | 0.00033 | 14.0 | 0.99 |
| 12 | 0.00097 | 11.0 | 0.85 |
| **Mean** | **0.00096** | **11.81** | **0.93** |
| **SEM** | **0.00014** | **1.25** | **0.01** |

**Table S2.**

Exponential demand curve parameters for individual subjects in Experiment 2. Note: the parameter k (range of consumption) is set to 2.2 log units.

| Paradigm | Group | Total DEGs | Up | Down |
|---|---|---|---|---|
| WIA | High (Resilient) vs. Sal | 1557 | 804 | 753 |
| | Low (Vulnerable) vs. Sal | 313 | 147 | 166 |
| Demand | High α (Resilient) vs. Sal | 223 | 91 | 132 |
| | Low α (Vulnerable) vs. Sal | 472 | 259 | 213 |
| Reinstatement | Low (Resilient) vs. Sal | 448 | 227 | 221 |
| | High (Vulnerable) vs. Sal | 488 | 278 | 210 |

**Table S3.**

Numbers of DEGs in each comparison group in males. log2(Fold Change) > 0.32, $p < 0.05$.

| Paradigm | Group | Total DEGs | Up | Down |
|---|---|---|---|---|
| WIA | High (Resilient) vs. Sal | 560 | 125 | 435 |
| | Low (Vulnerable) vs. Sal | 655 | 199 | 456 |
| Demand | High α (Resilient) vs. Sal | 304 | 104 | 200 |
| | Low α (Vulnerable) vs. Sal | 479 | 162 | 317 |
| Reinstatement | Low (Resilient) vs. Sal | 358 | 136 | 222 |
| | High (Vulnerable) vs. Sal | 301 | 155 | 146 |

**Table S4.**

Numbers of DEGs in each comparison group in females. log2(Fold Change) > 0.32, $p < 0.05$.

| Paradigm | Group | Total Diff. Sites | Up | Down |
|---|---|---|---|---|
| WIA | High (Resilient) vs. Sal | 1407 | 579 | 828 |
| | Low (Vulnerable) vs. Sal | 2312 | 681 | 1631 |
| | Res vs. Vul | 1766 | 1551 | 215 |
| Demand | High α (Resilient) vs. Sal | 2363 | 1319 | 1044 |
| | Low α (Vulnerable) vs. Sal | 3706 | 2555 | 1151 |
| | Res vs. Vul | 4631 | 1858 | 2773 |
| Reinstatement | Low (Resilient) vs. Sal | 3080 | 2430 | 650 |
| | High (Vulnerable) vs. Sal | 815 | 480 | 335 |
| | Res vs. Vul | 1873 | 1307 | 566 |

**Table S5.**
Numbers of differential sites in each comparison group in males. log2(Fold Change) > 0.32, padj < 0.05.

| Paradigm | Group | Total Diff. Sites | Up | Down |
|---|---|---|---|---|
| WIA | High (Resilient) vs. Sal | 861 | 369 | 492 |
| | Low (Vulnerable) vs. Sal | 1214 | 903 | 311 |
| | Res vs. Vul | 3915 | 1308 | 2607 |
| Demand | High α (Resilient) vs. Sal | 2581 | 1753 | 829 |
| | Low α (Vulnerable) vs. Sal | 9049 | 6078 | 2972 |
| | Res vs. Vul | 1273 | 523 | 750 |
| Reinstatement | Low (Resilient) vs. Sal | 613 | 177 | 436 |
| | High (Vulnerable) vs. Sal | 329 | 170 | 159 |
| | Res vs. Vul | 343 | 52 | 291 |

**Table S6.**

Numbers of differential sites in each comparison group in females. log2(Fold Change) > 0.32, padj < 0.05.

#### References

1. S. A. Roiko, A. C. Harris, M. G. LeSage, D. E. Keyler, P. R. Pentel, Passive immunization with a nicotine-specific monoclonal antibody decreases brain nicotine levels but does not precipitate withdrawal in nicotine-dependent rats. *Pharmacol Biochem Behav* **93**, 105 (2009).
2. A. C. Harris, C. Mattson, M. G. LeSage, D. E. Keyler, P. R. Pentel, Comparison of the behavioral effects of cigarette smoke and pure nicotine in rats. *Pharmacol Biochem Behav* **96**, 217–227 (2010).
3. A. C. Harris, D. Burroughs, P. R. Pentel, M. G. LeSage, Compensatory nicotine self-administration in rats during reduced access to nicotine: an animal model of smoking reduction. *Exp Clin Psychopharmacol* **16**, 86–97 (2008).
4. M. G. LeSage, D. E. Keyler, D. Shoeman, D. Raphael, G. Collins, P. R. Pentel, Continuous nicotine infusion reduces nicotine self-administration in rats with 23-h/day access to nicotine. *Pharmacol Biochem Behav* **72**, 279–289 (2002).
5. Y. Swain, P. Muelken, M. G. LeSage, J. C. Gewirtz, A. C. Harris, Locomotor activity does not predict individual differences in morphine self-administration in rats. *Pharmacol Biochem Behav* **166**, 48 (2018).
